# Tempo: an unsupervised Bayesian algorithm for circadian phase inference in single-cell transcriptomics

**DOI:** 10.1101/2022.03.15.484454

**Authors:** Benjamin J. Auerbach, Garret A. FitzGerald, Mingyao Li

## Abstract

The circadian clock is a 24-hour cellular timekeeping mechanism that temporally regulates human physiology. Answering several fundamental questions in circadian biology will require joint measures of single-cell circadian phases and transcriptomes. However, no widespread experimental approaches exist for this purpose. While computational approaches exist to infer cell phase directly from single-cell RNA-sequencing (scRNA-seq) data, existing methods yield poor circadian phase estimates, and do not quantify estimation uncertainty, which is essential for interpretation of results from highly sparse scRNA-seq data. To address these unmet needs, we developed Tempo, a Bayesian variational inference approach that incorporates domain knowledge of the clock and quantifies phase estimation uncertainty. Through simulations and analyses of real data, we demonstrate that Tempo yields more accurate estimates of circadian phase than existing methods and provides well-calibrated uncertainty quantifications. We further demonstrate that these properties generalize to the cell cycle. Tempo will facilitate large-scale studies of single-cell circadian transcription.

## Introduction

The circadian molecular clock is a 24-hour timekeeping mechanism found in nearly every cell in humans^1^. The time of the clock, referred to as circadian phase, is determined by the mRNA and protein concentrations of the clock’s constituent genes, referred to as clock or core clock genes^2^. Clock genes are organized in a transcriptional-translational feedback loop that enables cells to maintain self-sustained ∼24 hour oscillations in the concentrations of clock gene mRNA. Clock gene proteins additionally interact with cell-type-specific regulatory factors to drive rhythmic transcription of genes referred to as clock-controlled genes (CCGs). It is partially through these CCGs that circadian clocks generate rhythmic cellular behaviors, such as rhythms in hepatocyte glycogenesis^3^ and vascular smooth muscle cell contractability^4^. Though self-sustained, circadian clocks additionally rely on environmental cues, referred to as Zeitgebers (e.g. light), to update and optimize their timing via a process referred to as entrainment^5^.

Many open questions in chronobiology require single-cell resolution, such as the identification of cell-type-specific CCGs and the role of circadian phase in gating cell fate decisions. As droplet-based single-cell RNA-sequencing (scRNA-seq) measures genome-wide single-cell transcriptomes at high throughput, it has become an attractive tool to study many of these questions. Existing scRNA-seq studies of the clock have relied on time course designs^6, 7, 8^, in which cell clocks are presumed to be entrained by an external rhythmic stimulus, such as light. Assuming cell clocks are perfectly synchronized, sample timing over the cycle of the stimulus can be used as a direct proxy for the circadian phases of all cells in the sample. Nevertheless, this is a limiting assumption, as previous studies suggest cell circadian phases can differ by several hours within the same tissue *in vivo* and are determined by biological variables such as spatial location^9, 10, 11, 12^. Furthermore, chronobiologists may be interested in studying circadian transcriptional rhythms of cells in the absence of timing cues (e.g. unsynchronized cells in a dish). Breaking this assumption requires single-cell measures of circadian phase. One approach is to estimate cell circadian phases from gene expression directly, a task referred to as unsupervised phase inference.

Several algorithms have been developed for the similar task of unsupervised phase inference for cell cycle analysis using scRNA-seq data^13, 14, 15, 16^. However, the circadian cycle and cell cycle differ in two notable ways. First, while hundreds of “core” genes are known to oscillate over the cell cycle, many of which are highly expressed^17, 18^, the core circadian clock is only comprised of ∼20 moderately expressed genes^2^. Second, ∼100-1000 CCGs^6, 7, 8^ oscillate in a cell-type-specific manner over the circadian cycle and the identities of these genes are often unknown ahead of time. Due to the lower information content of clock genes and the challenge in identifying CCGs, existing unsupervised phase inference methods perform poorly when tasked with ordering cells over the circadian cycle. An optimal approach for estimating circadian phase in scRNA-seq should thus identify CCGs *de novo* and incorporate their information into phase estimates.

Existing unsupervised phase inference approaches were mainly developed for scRNA-seq data generated by plate-based approaches (e.g. Fluidigm C1). Relative to droplet-based techniques (e.g. 10X Genomics Chromium), plate-based approaches tend to capture fewer cells and more unique transcripts per cell^19^. As such, existing approaches have three key limitations when applied to droplet-based scRNA-seq data. First, existing approaches yield poor point estimates of cell phase due to transcript likelihood distribution choices that do not closely approximate the true generative distribution of droplet scRNA-seq. Second, existing approaches do not quantify the uncertainty of phase estimates. This becomes crucial for interpretation of results from highly sparse droplet scRNA-seq. Third, run times of existing approaches scale poorly with the number of cells, making analyzing droplet scRNA-seq untenable for many applications.

To address these unmet needs, we developed Tempo, a Bayesian variational inference approach, for circadian phase inference that works well for both droplet-based and plate-based scRNA-seq data. Tempo is fast, can incorporate domain knowledge, and yields uncertainty quantifications for the estimated circadian phases. Using both simulated data with ground-truths and real scRNA-seq data, we demonstrate Tempo’s ability to achieve state-of-the-art cell phase point estimates and well-calibrated cell phase uncertainty quantifications. Moreover, we demonstrate Tempo’s performance generalizes to other cyclical processes, such as the cell cycle.

## Results

### Overview of Tempo

Tempo assumes transcript counts of gene *j* in cell *i, X*_*ij*_, follow a Negative Binomial distribution. The mean proportion of transcript counts are presumed to follow a sinusoid. Thus, the mean of transcript counts of gene *j* in cell *i* is influenced by two factors: 1) gene-specific parameters, β_*j*_, describing the shape of the sinusoid and 2) the cell’s circadian phase, θ_*i*_. Given the observed data, *X*, and prior knowledge of all cell and gene parameters, *P*(θ, β), we seek the posterior distribution of the cell and gene parameters, *P*(θ, β|*X*). However, a closed-form solution for *P*(θ, β|*X*) is unknown and estimation using sampling techniques is computationally burdensome. For a computationally efficient solution, Tempo instead proposes an approximate posterior distribution *q*(θ, β) with differentiable parameters describing its shape. Tempo estimates the true posterior, *P*(θ, β|*X*), by maximizing its similarity with the approximate posterior, *q*(θ, β), through a two-step iterative process (Figure 1). As input, Tempo requires the observed data, *X*, prior knowledge *P*(θ, β), and a list of core clock genes. Tempo uses this information to initialize a list of cycling genes, which only includes the core clock genes at initialization, and the approximate posterior, *q*(θ, β). The approximate posterior *q*(θ, β) is formulated such that only cycling genes contribute information to the approximate posterior estimate of cell phases. In Step 1, Tempo optimizes *q*(θ, β) to minimize its KL divergence with *P*(θ, β|*X*) using only information from the current cycling genes. After this step, the marginal of *q*(θ, β) with respect to θ can be considered a rough estimate of the cell circadian phase posterior distributions based on only the current cycling genes. In Step 2, Tempo uses the cell phase posterior distributions from Step 1 to identify *de novo* cyclers. For the set of genes not currently identified as cyclers, approximate gene parameter distributions are fit, conditioned on the cell phase posterior distributions from Step 1. Tempo then selects *de novo* cycling genes as those best described by phase variation and adds them to the set of current cycling genes. Steps 1 and 2 are repeated until the core clock gene Bayesian evidence worsens or the maximum number of iterations is exceeded. The final result of the algorithm is the optimized approximate joint posterior distribution *q*(θ, β), which contains information about posterior cell phases and the set of identified *de novo* cycling genes.

**Figure 1:**
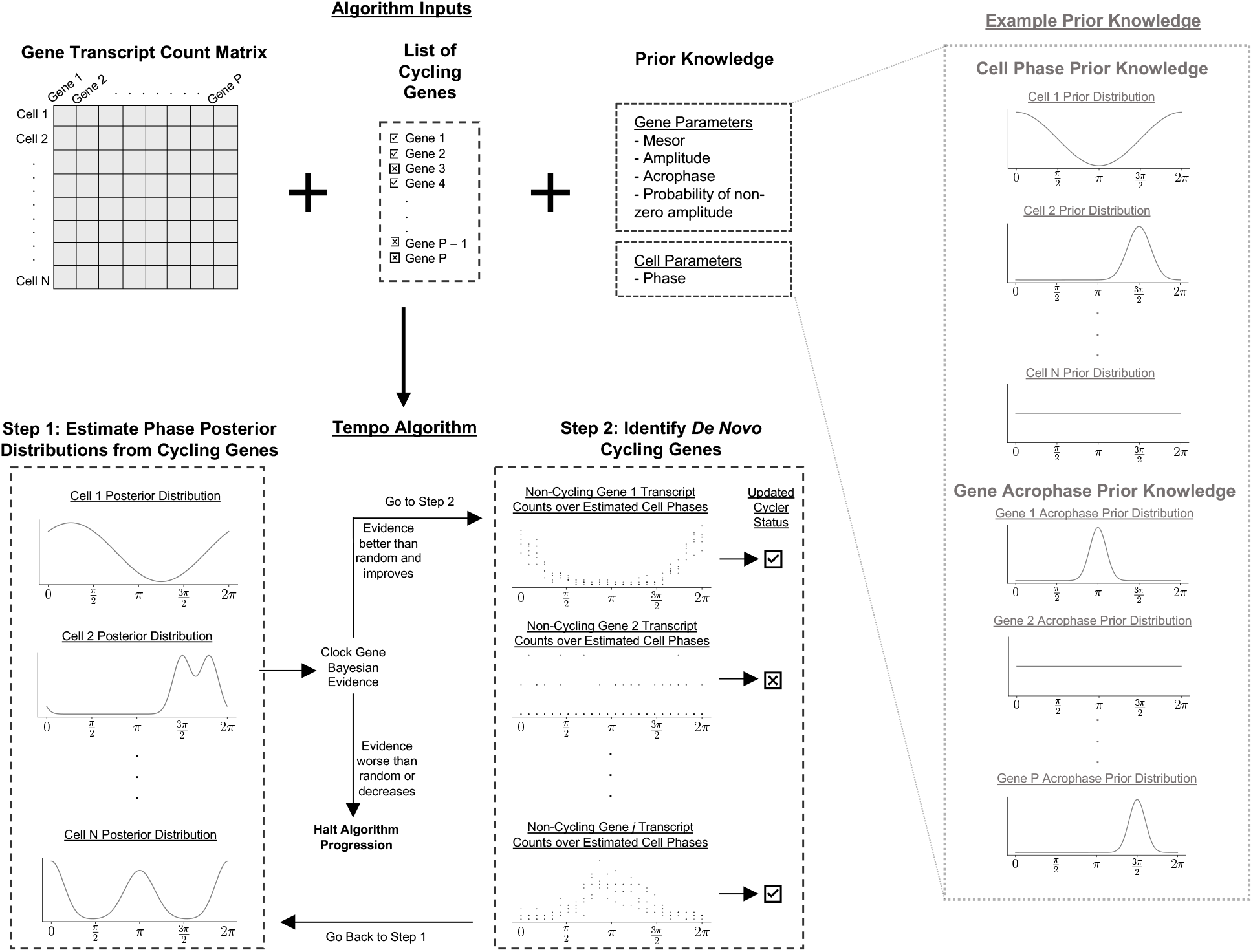
Tempo model overview. As input, users supply a cell transcript count matrix, list of cycling genes (e.g. circadian clock genes), and prior knowledge about the cell and gene parameters. Using the user-specified cycling genes, the count data and prior knowledge, in Step 1 Tempo computes approximate posterior distributions of the cell circadian phases. Using these, in Step 2 Tempo identifies *de novo* cycling genes with transcript counts that are well-explained by circadian phase variation. Tempo repeats Steps 1 and 2 until either the Bayesian evidence of the user-supplied cycling genes (e.g. circadian clock genes) worsens relative to previous iterations of the algorithm or is worse than random.

### Evaluations of Tempo on simulated data

We first assessed Tempo’s performance on simulated scRNA-seq data generated from Tempo’s Negative Binomial count model where sinusoidal gene parameters, including mesor, amplitude, and acrophase, were estimated from a light-dark cycle time course scRNA-seq dataset. This approach was used to simulate scRNA-seq datasets collected from either a single sample of unsynchronized cells or from cells sampled every 4 hours over a 24-hour light-dark cycle time course (i.e. ZT0, ZT6, ZT12, and ZT18). Details on the estimation of gene parameters and generative model used for simulations can be found in Methods.

Using these simulated scRNA-seq data, we first determined whether Tempo can accurately estimate circadian phase when considering only the core clock genes as input. Tempo was run using a non-informative prior over cell phases. To mimic informative but imperfect gene priors, core clock acrophase prior locations were shifted from their true values. Shifts were drawn from standard normal distributions, scaled by 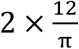 (i.e. standard deviation of 2 hours), and added to the true acrophase values to yield acrophase prior locations. The prior clock acrophase scales of the Von Mises distribution were set such that the width of the 95% interval surrounding the prior acrophase locations was 4 hours. Cell phase point estimate errors were visualized as empirical cumulative distribution functions (eCDFs). For both unsynchronized and time course datasets, and across a wide range of cell numbers (500 to 5000 cells) and library sizes (3000 to 20000 UMIs), Tempo yields point estimate error eCDFs slightly worse than the optimum (Figure 2a, Supplemental Figures 1-9a). The optimum was obtained by computing the maximum likelihood phases using the true generative model as the likelihood model and considering all true cycling genes as input and setting gene parameters to their true values. For comparison, we analyzed the simulated data using existing unsupervised phase inference approaches with run time characteristics suitable for droplet scRNA-seq, including Cyclops^20^, Cyclum^16^, and PCA^21^. These competing approaches yielded non-random performance, but demonstrated a stark decrease in performance for data with smaller library sizes.

**Figure 2:**
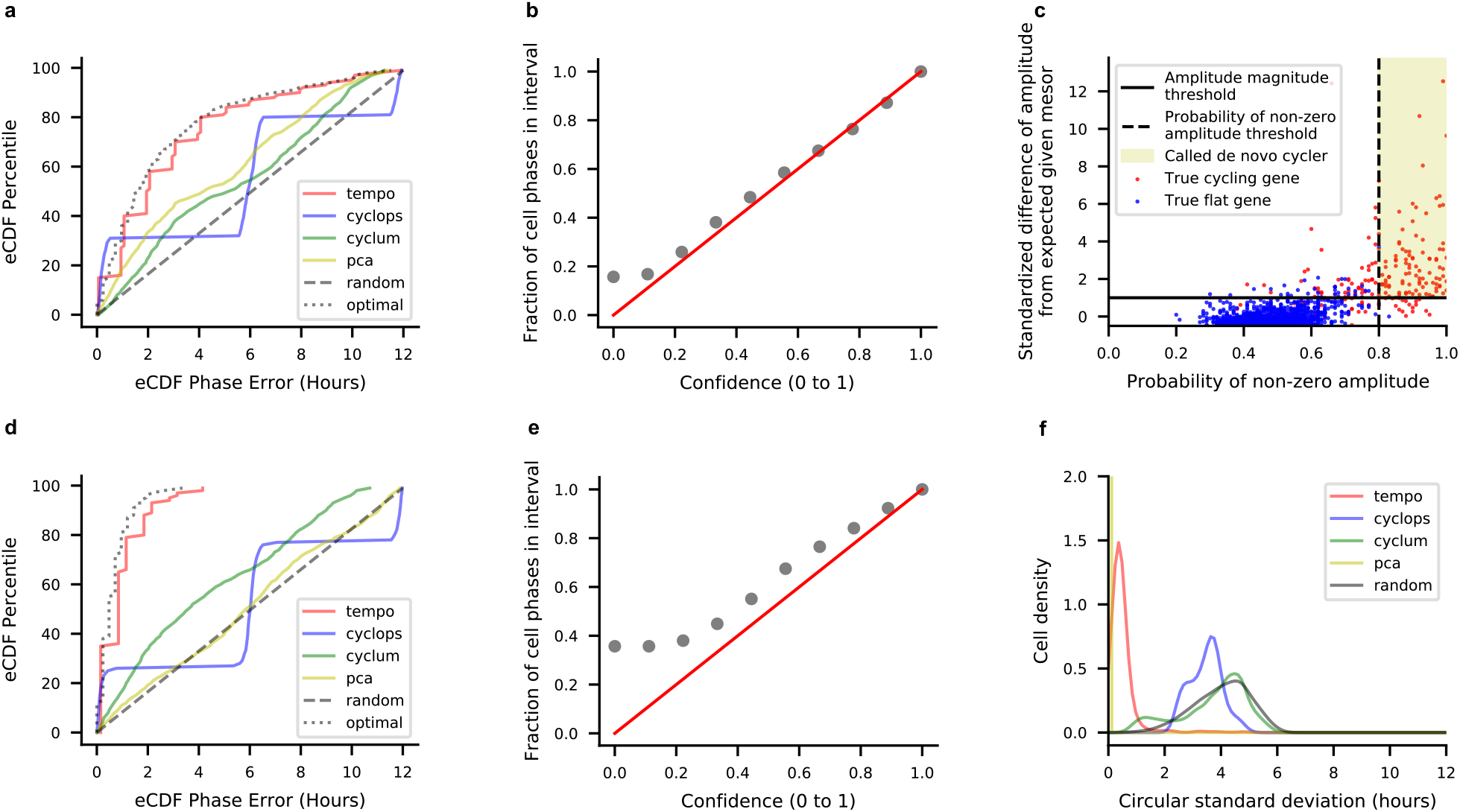
Results on a simulated scRNA-seq dataset of 1000 cells collected at CT0, CT6, CT12, and CT18 with mean library size of 10000 UMI. **a)** Empirical cumulative distribution function (eCDF) of the errors for each method’s cell phase point estimates, where all methods were run using the true core clock genes as input. **b)** Calibration of Tempo’s uncertainty estimates when run using the true core clock genes as input. **c)** Tempo’s *de novo* cycler detection procedure. The x-axis represents the *maximum a posteriori (*MAP) fraction of samples with non-zero amplitude for a given gene, and captures whether a gene is better described by sinusoidal or flat variation over the circadian cycle. The y-axis statistic measures deviation of a gene’s MAP amplitude from its expected MAP amplitude given its MAP mesor, reported in terms of a Pearson residual. Large positive values indicate a gene has a larger amplitude than expected given its mesor. Details of the Pearson residual computation can be viewed in Supplemental Information section 8. **d)** eCDF of the errors for method cell phase point estimates, where methods were run using default settings. **e)** Calibration of Tempo’s uncertainty estimates when run with default settings. **f)** Method stability analysis. Methods were run 5 times (with default settings) on the dataset. The circular standard deviation of predictions for each cell was computed and visualized as a distribution.

We further probed whether the uncertainty quantifications associated with Tempo’s phase estimates using the core clock genes alone were well-calibrated. We assessed the relationship between the confidence of the approximate posterior’s credible interval and the corresponding fraction of intervals containing the true cell phase. Credible intervals were computed using the Highest Density Region approach^22^. Encouragingly, this analysis (Figure 2b, Supplemental Figures 1-9b) suggests Tempo’s uncertainty quantifications are well-calibrated for both unsynchronized and time course data. Tempo’s credible intervals are slightly conservative, which reflects the propagation of uncertainty from the gene parameters.

Given that Tempo can estimate cell phase from the core clock genes alone, we next assessed the feasibility of *de novo* cycler detection and the potential use of *de novo* cyclers to improve cell phase point estimates. Tempo was run on simulated datasets considering all genes as input and using *de novo* cycler detection. For comparison, Cyclops, Cyclum, and PCA were run with default settings considering all genes as input. For both unsynchronized and time course datasets, and across a range of simulation settings, Tempo identifies *de novo* cycling genes with high specificity and sensitivity (Figure 2c, Supplemental Figures 1-9c). Notably, incorporating *de novo* cyclers with core clock genes improves cell phase point estimates (Figure 2d, Supplemental Figures 1-9d). In comparison, competing methods did not see a significant improvement in point estimates when considering all genes as input. Phase uncertainties remained well-calibrated, albeit more conservative, when incorporating *de novo* cyclers (Figure 2e, Supplemental Figures 1-9e). This suggests that *de novo* cycler detection can be a valuable tool for circadian phase estimation.

We further assessed the stability of Tempo’s predictions. Tempo’s results are stochastic due to the sampling required to compute the objective function for both cell phase estimation and *de novo* cycler detection. It is crucial to validate that the default number of samples used to compute the objective function yields stable results. To assess method stability, methods were run on the same simulated dataset multiple times. For each method, the circular standard deviation (reported in hours) of all cells was computed and visualized as a distribution. For comparison, circular standard deviation distributions were also computed by randomly drawing cell phases from a circular uniform distribution. Tempo’s median circular standard deviation was less than 1 hour for all simulation settings for which stability was evaluated (Figure 2f, Supplemental Figures 1f, 2f, 7f). Of note, Cyclum and Cyclops yielded highly unstable results for the simulation settings evaluated.

### Tempo accurately estimates circadian phase from time course data

Encouraged by Tempo’s performance on simulated data, we next assessed the quality of Tempo’s circadian phase estimates on real droplet-based scRNA-seq data. We generated a deeply sequenced scRNA-seq dataset using the 10X Genomics Chromium platform from mouse aorta collected every 4 hours (i.e. ZT0, ZT6, ZT12, and ZT18) over a 24-hour light-dark cycle. This high-quality dataset yielded 18863 vascular smooth muscle cells (SMCs), 3135 fibroblasts, 288 macrophages, and 287 endothelial cells with median library sizes of 13646, 7412, 6846.5, and 7389 unique molecular identifiers (UMIs), respectively. To benchmark Tempo on this dataset, we compared its performance with Cyclops, Cyclum, and PCA. All methods were run on each cell type twice. First, methods were run using default settings considering all genes as input. Second, methods were run by restricting the input genes to only the core clock genes. Tempo was run using a noninformative prior over cell phases. Additional detail on the initialization of the core clock gene priors is provided in Methods. The resulting cell phase predictions for each individual time point can be viewed in Supplemental Figures 10-14. For each cell type, we first compared the circadian phase point estimates of individual cells to their sample collection phase in the light-dark cycle. The distribution of the difference between the two phases over all cells was visualized as an eCDF (Figure 3a and Supplemental Figures 15-21a). Tempo favored cell phase estimates based on the core clock genes alone rather than those including *de novo* cyclers for all real light-dark cycle cell type datasets. For all cell types Tempo’s point estimates demonstrate a substantial improvement over the alternative approaches we analyzed (Supplementary Tables 1 and 2). Moreover, on these data Tempo demonstrates well-calibrated uncertainties (Figure 3b and Supplemental Figures 15-21b), suggesting its uncertainty quantification is meaningful and can aid interpretation of results. Methods were also applied to time course droplet scRNA-seq of 18378 mouse hepatocytes from *Droin et al*.^*6*^. On these sparser data (median library size of 1965 UMI), Tempo again demonstrates improved point estimates over competing methods and well-calibrated uncertainties (Supplementary Figures 22-23 a and b).

**Figure 3:**
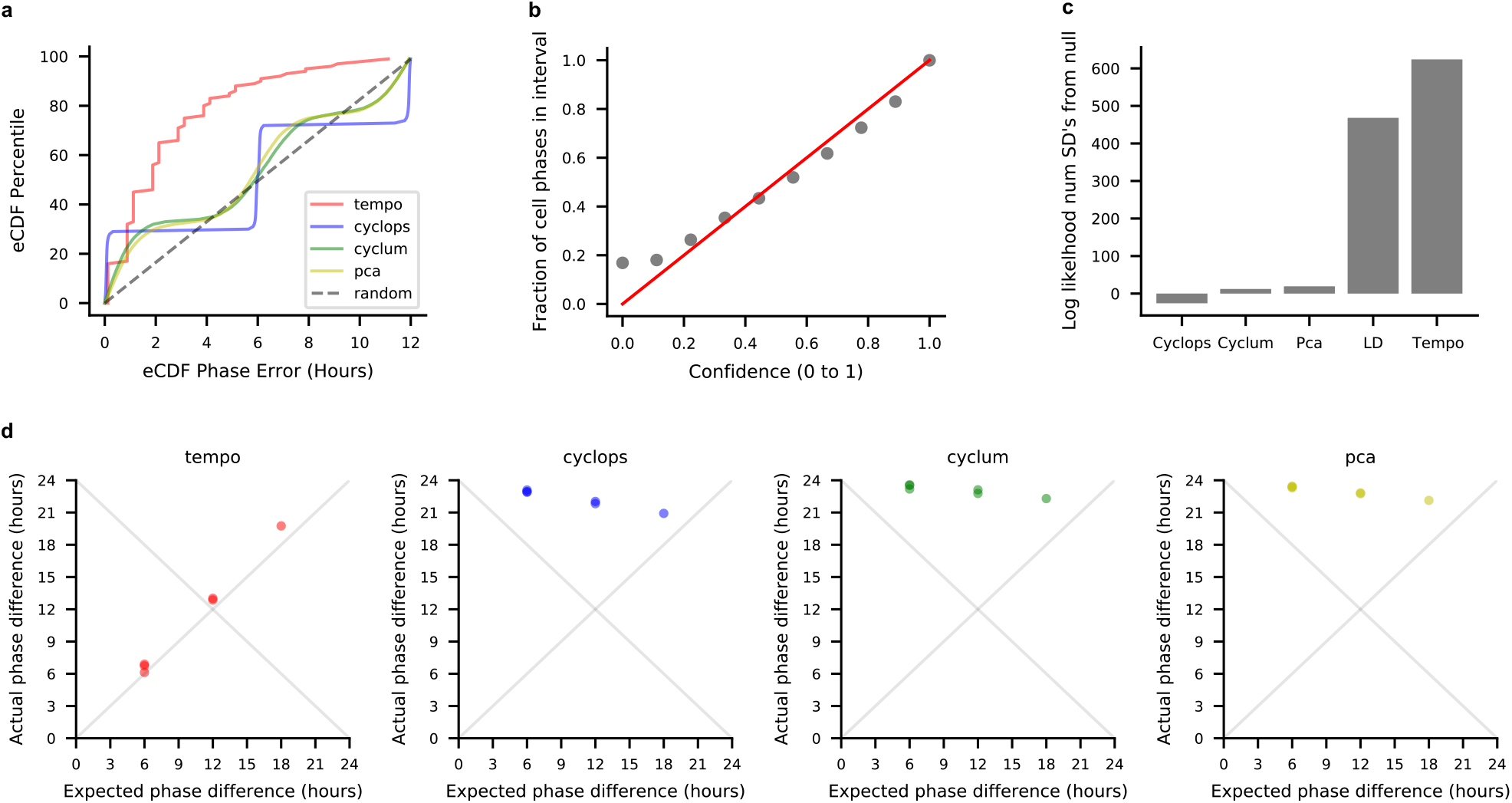
Method results (default settings) on light-dark cycle aorta smooth muscle cells. Treating the sample collection phase in the light-dark cycle as the true cell circadian phases: **a)** eCDF of the errors for each method’s cell phase point estimates **b)** Calibration of Tempo’s uncertainty estimates. **c)** Method out of sample core clock gene likelihood analysis. LD corresponds to treating sample collection times as the true cell phases. Out of sample core clock likelihoods were computed for each method, and reported in terms of standard deviations from the median of a distribution of likelihoods associated with phase assignments drawn from a random uniform distribution. **d)** Method relative shift analysis. Each dot represents a pair of sample collection times in the light-dark cycle (e.g. all 6 possible pairs of ZT0, ZT6, ZT12, ZT18), and conveys the relationship between the expected phase difference between a pair of time points and the actual phase difference for each method. As the phase difference is a circular random variable, methods with points lying along either y = x or y = 24 – x denote perfect performance.

The above evaluations assume each cell’s circadian phase is equal to its sample collection phase in the light-dark cycle. While we anticipate the true cell phase is close to the sample collection phase in young, healthy mice, the collection phase may not exactly equal the true circadian phase of individual cells if they are imperfectly synchronized. As such, methods were evaluated on light-dark cycle data according to two additional criteria independent of single-cell phase ground truths. First, the difference between cell phase estimate distributions for any pair of sample collection phases should equal the difference in the collection phases. Based on this rationale, we calculated the expected relative shift in phase distributions across pairs of sample collection times and their respective cells. As shown in Figure 3d and Supplemental Figures 15-17d,18-23c, Tempo recapitulated the expected relative shift for all cell types. Alternative approaches often did not capture the anticipated relative phase shift between sample collection times, regardless of whether they were run with the core clock genes or the full gene set. Second, methods with phase estimates closest to the truth should best explain core clock expression, in terms of likelihood. Moreover, phase estimate parameters should generalize to explaining the expression of clock gene transcripts in unseen cells. As such, methods were evaluated for their ability to explain core clock expression on a holdout set of cells, as measured by the likelihood of core clock expression using the method’s cell phase estimates (Figure 3c and Supplemental Figures 15-17c). Details of the computation of holdout cell core clock expression likelihood can be viewed in Methods. Tempo explained holdout clock expression the best across all cell types. Intriguingly, for the aorta SMCs and fibroblasts, Tempo explains core clock expression better than sample collection phase. This suggests Tempo may meaningfully identify out-of-phase cells collected over light-dark cycles.

### Tempo’s performance generalizes to other cyclical processes and scRNA-seq platforms

To determine whether Tempo’s performance generalizes to other cyclical processes and scRNA-seq platforms, we applied it to infer cell cycle phase from scRNA-seq data of human induced pluripotent stem cells (iPSCs) generated by *Hsiao et al*.^*23*^ using the Fluidigm C1 platform. This dataset also contains continuous cell phase labels measured by two fluorescent cell cycle reporters for each cell, which are treated as the ground truth cell phases when evaluating the performance of each method. We benchmarked Tempo with Cyclops, Cyclum, and PCA. All methods were run on each cell type twice. First, methods were run using default settings considering all genes as input. Second, methods were run by restricting the input genes to a filtered list of annotated cell cycle genes from CycleBase^17^ (details of which can be viewed in Methods). Tempo was run using a noninformative prior over cell phases. The distribution of the cell phase errors was visualized as an eCDF (Figure 4a). Similar to Tempo’s circadian phase inference results on the light-dark cycle data, Tempo favored cell phase estimates based on the annotated cell cycle genes alone rather than those including *de novo* cyclers. Although not principally developed for cell cycle phase inference, Tempo yielded comparable results to existing approaches benchmarked for cell cycle phase inference, such as Cyclum. Tempo had a median point estimate phase error of 4.35 hours, compared to 5.02, 4.36, and 3.83 for Cyclops, Cyclum, and PCA. While obtaining similarly performing point estimates to existing standards, Tempo additionally yielded well-calibrated, albeit slightly overconfident, uncertainty quantifications on these data (Figure 4b). To lend confidence in the robustness of these cell cycle results, we further assessed methods on scRNA-seq data from 288 mouse embryonic stem cells (mESCs) from *Buettner et al*.^24^ that were discretely assigned to G1, S, and G2M via flow sorting. On these data, Tempo’s point estimates yield comparable results to competing methods and well-calibrated uncertainties (Figure 4c,d).

**Figure 4:**
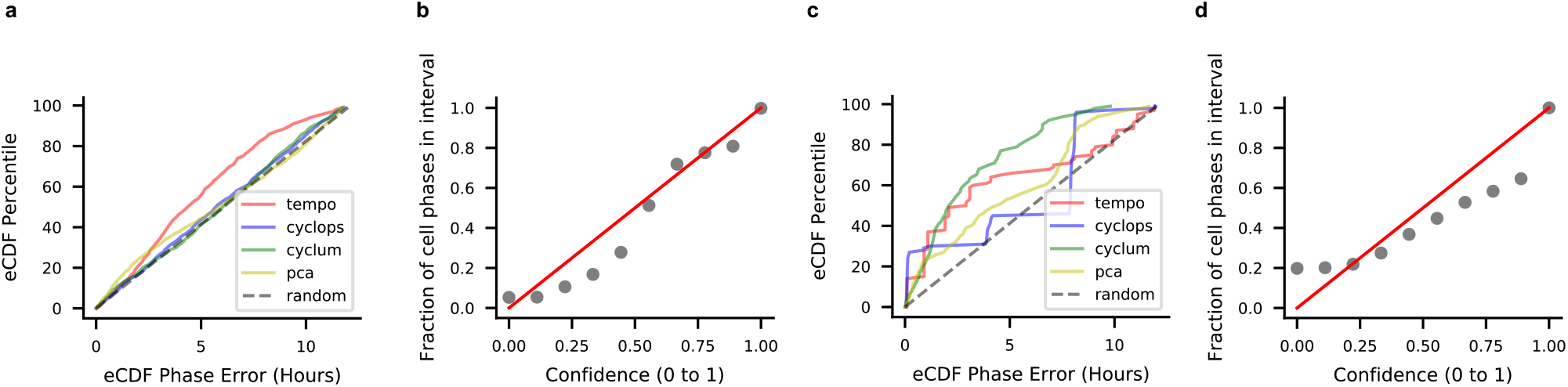
Assessing method cell cycle phase estimates on cells from *Hsiao et al*. scRNA-seq: **a)** eCDF of the errors for method cell phase point estimates, where all methods are run with default settings. **b)** Calibration of Tempo’s uncertainty estimates when run with default settings. Assessing method cell cycle phase estimates on cells from *Buettner et al*. scRNA-seq: **c)** eCDF of the errors for method cell phase point estimates, where all methods are run with default settings. **d)** Calibration of Tempo’s uncertainty estimates when run with default settings.

## Discussion

Single-cell transcriptomics offers an unprecedented opportunity to improve our understanding of circadian transcription. Nonetheless, its impact has been limited by the assumption that sample collection times reflect cell circadian phases. In lieu of widespread experimental approaches that jointly measure single-cell phase and transcriptomes, computational phase inference tools can be applied to estimate cell phases from scRNA-seq directly. However, existing tools yield poor phase estimates and do not quantify estimation uncertainty.

To address these challenges, we developed Tempo, a Bayesian algorithm for single-cell phase inference from scRNA-seq data. Through a combination of both simulated and real data analyses, we demonstrate that Tempo yields state-of-the-art point estimates. Moreover, Tempo empowers better phase estimate interpretation through well-calibrated uncertainty quantifications and measurement of improvement over random phase assignments. While initially developed for circadian phase inference, we further demonstrate Tempo’s performance generalizes to other cyclical processes, such as the cell cycle. Lastly, Tempo’s run time characteristics are amenable to larger droplet-based scRNA-seq datasets. Across all datasets analyzed, Tempo completed within 1 hour using a desktop CPU with a 3.2GHz Intel Xeon W processor and 32GB RAM.

Given Tempo’s performance, we are encouraged by its immediate potential for use in circadian research. More specifically, our tool may make it possible to characterize circadian transcription parameters using single samples of unsynchronized cell cultures. This experimental paradigm can enable cost-effective circadian transcription studies of human subjects and be used to study the role of cell-cell interactions on circadian transcription. Further, our tool can help researchers study circadian phase heterogeneity and its biological determinants in tissue contexts (e.g. spatial location) from time course data. Indeed, our analyses suggest Tempo meaningfully identifies out-of-phase cells in mouse aorta (Figure 3c, Supplemental Figures 15-17c). Lastly, cell populations demonstrate cell-cell heterogeneity in circadian gene expression parameters, such as acrophase and amplitude of clock genes etc. This Bayesian approach naturally captures this biological variation and facilitate its interpretation.

Although Tempo has achieved several advances in unsupervised phase inference, opportunities for future improvements exist. First, *de novo* cyclers do not improve point estimates in real scRNA-seq datasets. While incorporating *de novo* cyclers improves point estimates in simulated data, *de novo* cyclers decreased evidence of core clock expression in the real scRNA-seq datasets we analyzed. This may be due, in part, to the assumption that the expression means of CCGs follow sinusoidal patterns across the circadian cycle. Future efforts might rely on approaches that can model more flexible CCG waveforms. Second, our method does not explicitly model the contribution of technical effects to expression variation. While less important for applications to single-sample unsynchronized scRNA-seq data, this becomes more necessary for data collected as multiple samples over time.

While we await widespread experimental approaches to pair single-cell clock reporters with transcriptomics, it is important to keep in mind that clock reporters with even zero measurement error will contain phase uncertainty due to the inherent stochasticity of the clock. Thus, single-cell reporter phases may be best used as prior knowledge to unsupervised phase inference algorithms, such as Tempo.

In summary, we developed Tempo, a Bayesian algorithm for circadian phase inference from scRNA-seq data. Tempo yields state-of the-art point estimates of circadian phase, and well-calibrated uncertainties. These properties generalize to the cell cycle. Well-calibrated uncertainties will enable investigators to understand the robustness of cell phase estimates made from highly sparse scRNA-seq data and to account for uncertainty in downstream analyses. Further, the quality of Tempo’s phase estimates may make it possible to identify out-of-phase cells from tissue collected over time courses and to estimate circadian parameters from unsynchronized cell populations using scRNA-seq.

## Acknowledgements

This work was supported by 2U54TR001878 (G.A.F), R01GM125301 (M.L.), and 2T32HG000046-21 (B.J.A.). We thank Sean Anderson, Soon Yew Tang, and Ronan Lordan for generating the scRNA-seq data from mouse aorta, and Christopher J. Adams for a critical reading of the manuscript and testing of the software package.

## Author contributions

This study was conceived of by B.J.A. and led by B.J.A. and M.L.. B.J.A. designed the model and algorithm. B.J.A. implemented the Tempo software and led data analysis with input from M.L. and G.A.F.. B.J.A. wrote the paper with feedback from M.L. and G.A.F..

## Software availability

Tempo is provided as an open-source software package available at https://github.com/bauerbach95/tempo.

## Methods

### Tempo model

#### Likelihood model

As input, Tempo requires an *n* × *p* transcript count matrix *X*, where *n* is the number of cells and *p* is the number of genes. For gene *j* in cell *i*, the unique molecular identifier (UMI) count *X*_*ij*_ is assumed to follow a Negative Binomial (NB) distribution. The expected log proportion, log *λ*_*ij*_, of gene *j*’s transcripts in cell *i* is defined by a sinusoid with 4 parameters: 1) the mesor, *μ*_*j*_, which controls the mean of a gene’s proportion over the circadian cycle, 2) the amplitude, *A*_*j*_, or the maximum deviation of the gene’s proportion from the mesor over the circadian cycle, 3) the acrophase, *ϕ*_*j*_, or the peak time of the gene’s proportion over the circadian cycle, and 4) an indicator, *Q*_*j*_, describing whether the gene has non-zero amplitude. These sinusoidal gene parameters are assumed to be shared across all cells. As such, observed expression differences across cells are explained by differences in 1) latent cell phase, *θ*_*i*_, 2) cell library size, *L*_*i*_, and 3) random variation described by the Negative Binomial distribution.

The distribution of *X*_*ij*_ is modeled as

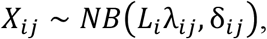

where

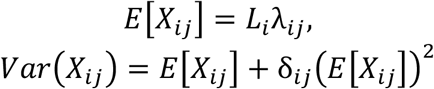

and

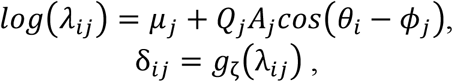

wher*e g*_*ζ*_(λ_*ij*_)is a deterministic polynomial function parameterized by *ζ* (shared by all cells and genes) describing the relationship between transcript proportion λ_*ij*_ and the dispersion δ_*ij*_. Details on the estimation of ζ can be found in Supplemental Information section 1.

#### Prior knowledge of gene and cell parameters

Prior knowledge may be known about cell phases (e.g. based on single-cell clock gene reporters or cell collection time); in this case, users can specify prior knowledge about the phase of cell *i* as a Von Mises distribution (a circular analog to the normal distribution),

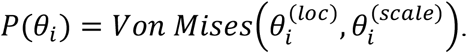

In the absence of prior knowledge about cell phases, Tempo uses a non-informative Hyperspherical Uniform prior for each cell phase by default.

For the gene parameters, prior knowledge about the mesor of gene *j* can be specified as a normal distribution:

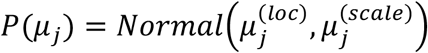

In practice, we use an empirical Bayesian approach to set 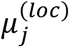 equal to the log proportion of transcripts for each gene.

Prior knowledge about the acrophase of gene *j* may exist, (e.g. from bulk circadian transcriptomics data), in which case prior knowledge can be specified in terms of a Von Mises distribution. In the case of the core circadian clock genes, prior knowledge is typically known. Otherwise, by default Tempo assumes a non-informative Hyperspherical Uniform prior.

The algorithm additionally requires the user to specify a reference gene, the peak time of which defines the start of the circadian cycle. To enforce this, the prior acrophase distribution of the reference gene is set to a point mass centered at 0 radians. By default, the algorithm uses the core circadian clock gene, Arntl, as the reference gene defining the start of the cycle.

Prior knowledge about the amplitude of gene *j* is specified as a transformed Beta distribution:

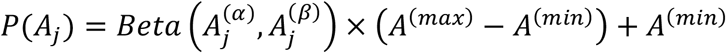

where *A*^(*min*)^ and *A*^(*max*)^ denote the minimum and maximum possible amplitude (which is shared by all genes). By default, Tempo sets 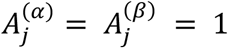, which assumes a non-informative prior over the domain of possible amplitude values.

Prior knowledge about whether a gene *j* has non-zero amplitude is specified in terms of a hierarchical Beta-Bernoulli:

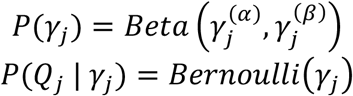

where samples of *γ*_*j*_ denote the success probability of a gene having non-zero amplitude and the user specifies the shape parameters of the Beta distribution. For genes not part of the user-specified list of core clock genes, Tempo sets 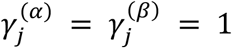, by default.

#### Approximate posterior inference

Using our prior knowledge of the cell and gene parameters and the observed data, we seek the following joint posterior distribution of our cell and gene parameters:

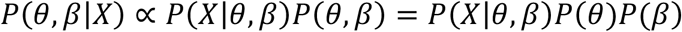

where *θ* is an *n*-dimensional vector containing the phase for each cell, *β* is a *p-*dimensional vector containing the parameters for each gene (i.e. β_*j*_ = (µ_*j*_, *A*_*j*_, ϕ_*j*_, *Q*_*j*_, γ_*j*_)), and

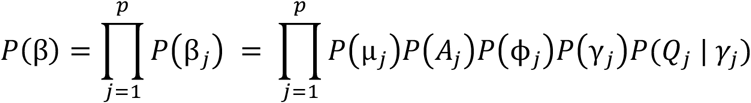

and

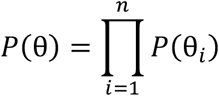

No known analytic solution exists to *P*(*θ, β*|*X*). Moreover, asymptotically exact estimation approaches, such as Markov-chain Monte Carlo and full grid sampling, do not scale well to droplet-based scRNA-seq datasets that can contain thousands (and sometimes tens of thousands) of cells.

For a computationally efficient solution to estimate *P*(*θ, β*|*X*), Tempo uses variational inference, an optimization-based approximate Bayesian inference approach. In brief, Tempo uses the list of core clock genes and prior knowledge to initialize an approximate joint posterior distribution *q*(θ, β) with differentiable parameters describing its shape. *q*(*θ, β*) is parameterized such that it includes a list of cycling genes and only cycling genes contribute information to the estimate of θ. At initialization, the cycling gene list only includes the user-supplied core clock genes. Tempo optimizes *q*(*θ, β*) to approximate *P*(*θ, β*|*X*) by minimizing their KL divergence through an iterative two-step procedure. In Step 1, Tempo estimates the cell phase distributions and gene parameter distributions using only information from the current cycling genes to minimize the KL divergence between the true joint posterior distribution and the approximate joint posterior distribution. In Step 2, Tempo identifies *de novo* cycling genes whose expression variation is well-described by circadian variation. Approximate gene parameter distributions are computed for current non-cycling genes conditional on the cell phase distributions computed in Step 1 and conditional on the non-cycling genes having non-zero amplitude (i.e. *Q*_*j*_ is set to 1). Tempo then identifies non-cycling genes with expression that is sufficiently better explained by sinusoidal than flat variation and genes with sufficiently high amplitude as *de novo* cycling genes. *De novo* cyclers are then added to the cycling gene list. Steps 1 and 2 are repeated, estimating the cell phase posterior distribution from the current cycling genes and identifying *de novo* cyclers, until the algorithm’s stopping criteria are met. The final results of the algorithm are a set of identified cycling genes (the core clock genes and identified *de novo* cycling genes), and the approximate posterior distributions of all gene and cell parameters. Additional details of the initialization, generative process, and iterative two-step optimization procedure of *q*(θ, β) can be viewed in Supplemental Information sections 2 and 3.

### Generation of simulated scRNA-seq data

Realistic gene expression parameters were first estimated from the aorta SMC scRNA-seq dataset. Treating the light-dark sample collection time as the true cell phases, gene posterior distributions were estimated using variational inference according to Tempo’s likelihood model. Genes with high amplitudes (Pearson residuals greater than 2 for difference between MAP amplitude and expected MAP amplitude conditional on MAP mesor) and cycler probability point estimates greater than 0.95 were called as cycling genes, in addition to the annotated core clock genes.

Using the gene parameters estimated from real scRNA-seq, data were simulated according to the following parameters: 1) the number of cells; 2) the mean and standard deviation of log library size of the cells; 3) the number of equally spaced cell phases; 4) the number of flat genes; 5) the number of CCGs. Log library sizes were drawn from a normal distribution. Discrete cell phases were drawn from a multinomial distribution with uniform probabilities. Flat genes and CCGs, and their respective sinusoidal parameters, were sampled with replacement. Given these parameters, scRNA-seq datasets were generated by sampling from our model generative distribution. To simulate scRNA-seq data of unsynchronized cell cultures, cell phases were drawn from 23 equally spaced phases over the circadian cycle. To simulate light-dark cycle time courses, cell phases were drawn from 4 equally spaced phases over the circadian cycle.

### Optimal phase shifting procedure and computation of phase estimation errors

Given the phase estimate of cell *i* made by method *m*, 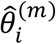, we would like to compute its error relative to the true cell phase, *θ*_*i*_. However, computing the phase error of method estimates is not straightforward. Cyclops, Cyclum, and PCA do not use information about which gene’s peak defines the start of the circadian cycle. As such, the absolute latent cell phase estimates of these methods are arbitrary, though the relative ordering of the cells is not. To deal with this, each method *m*’s phase estimates are shifted by *s*^*(m)*^ that produces the minimum total error over all cells. *s*^*(m)*^ is computed as:

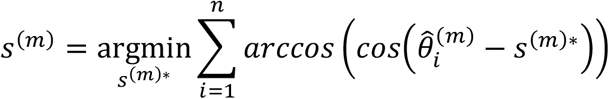

The method phase prediction for each cell *i*, 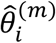, is then shifted:

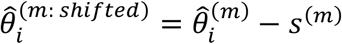

Given the true cells phases, θ, and method-specific shifted phase estimates, 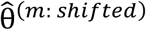, we compute the error (in hours) of the phase point estimate made by method *m* for cell *i*, 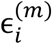:

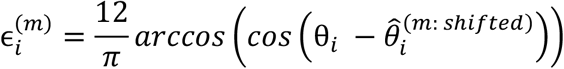

### Simulated data circadian phase estimation

The acrophase priors used by Tempo for the simulated core clock genes were set as follows. First, the prior acrophase location was shifted from the true acrophase value. This was done by drawing a shift from a standard normal distribution, scaling the shift by 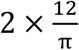, and then adding the shift to the true acrophase value. Second, the prior acrophase scale of the Von Mises distribution was set such that the width of the 95% interval surrounding the prior acrophase location was 4 hours.

Tempo, Cyclops, Cyclum, and PCA were run two times using the algorithms’ default settings: first, considering all genes, and, second, restricting the data to the core clock genes.

### Simulated data model stability analysis

For the simulated datasets with results shown in Figure 2 and Supplemental Figures 1, 2, and 7, Tempo, Cyclops, Cyclum, and PCA were each run 5 times. As a baseline method to compare against, 5 random cell phase predictions were also made for each cell by drawing from a uniform distribution over [0, 2π]. Predictions from Tempo, Cyclops, Cyclum, PCA, and the random method yielded a matrix 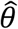, where 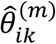 denotes the phase point estimate for cell *i* method *m* and run *k*. For each cell – method pair, the expected phase across runs for each cell, 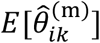, was computed using the SciPy^25^ circular mean function. The stability (reported in hours) of method *j*’s predictions for cell *i*, 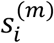, was then computed as follows:

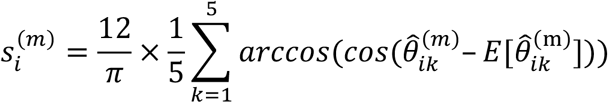

Smoothed distributions (Gaussian kernel with bandwidth 0.2 for non-PCA methods, 0.01 for PCA) of this stability metric can viewed in Figures 2f and Supplemental Figures 1f, 2f, 7f.

### Aorta data collection and data processing

8-week-old C57BL6 male mice were entrained to a 12:12 light-dark cycle for 2 weeks. At ZT0, ZT6, ZT12, or ZT18 mice were sacrificed (2 per time point). For each mouse, whole aorta was dissected, and cells were disassociated. Cells from mice sacrificed at the same time point were pooled to form single-cell suspensions (4 suspensions in total). Single cells were then captured and barcoded using the 10X Genomics Chromium platform and sequenced using an Illumina NovaSeq S2 flow cell.

Cell barcode detection, read alignment, and transcript quantification were performed using the 10X Genomics Cell Ranger pipeline. Cells with unique molecular identifier (UMI) count less than 1000 and mitochondrial fraction greater than 0.2 were discarded. In addition, cell doublets were detected by the Scrublet program^26^ and discarded. A low dimensional representation for the cells was obtained using UMAP^27^ using z-score log1p library size normalized counts of the aorta cell type markers from *Pan et al*.^*28*^. Using the UMAP representation as input, cells were then clustered using the ScanPy^29^ implementation of the Leiden algorithm^30^. This yielded 5 clusters corresponding to vascular smooth muscle cells, fibroblasts, macrophages, endothelial cells, and T cells. As only 34 T cells were detected, they were not included in the analyses detailed in this manuscript.

### Light-dark cycle data circadian phase estimation

The acrophase priors used by Tempo for the core clock genes were set as follows. For the aorta cell types, prior acrophase locations were set to the estimated acrophases in bulk liver RNA-seq from *Zhang et al*.^*31*^. For the hepatocytes from *Droin et al*.^*6*^, prior acrophase locations were set to the estimated acrophases in bulk liver RNA-seq from *Zhang et al*. The width of the prior acrophase 95% intervals were set to 2 hours for both the aorta and hepatocyte data. Arntl was treated as the reference gene.

### Light-dark cycle data out of sample clock likelihood analysis

Out of sample clock likelihoods were computed for each method for the aorta SMCs (5000 train and 13863 test cells) and aorta fibroblasts (1500 train and 1635 test cells). For a given method *m* with training set cell phase point estimates 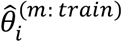 and corresponding training set gene parameter point estimates β^(*m*:*train*)^, we compute the test set core clock expression likelihood *D*^(*m*)^ as follows:

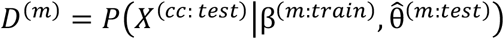

where 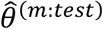 are the test set cell phase point estimates from method *m*, and *X*^(*cc*: *test*)^ is the test set core clock transcript count matrix generated according to:

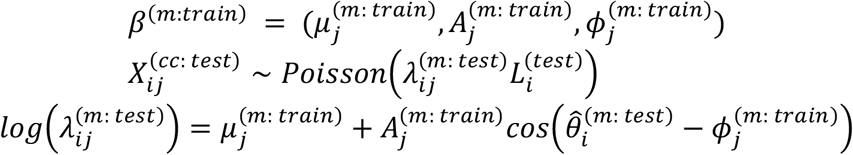

For Tempo, *β*^(*m*:*train*)^is computed during training. Cyclops, Cyclum, and PCA do not explicitly estimate *β*^(*m*:*train*)^ during training, but instead learn a mapping 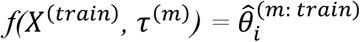, where *τ*^(*m*)^ represents the learned parameters of method *m* on the training data. For Cyclops, Cyclum, and PCA, we find the point estimate *β*^(*m*:*train*)^ that maximizes the expression likelihood under the following Poisson GLM :

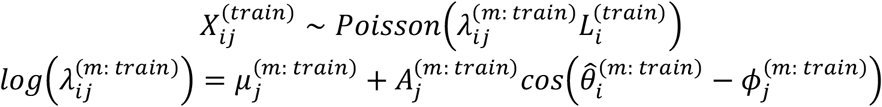

As a lower bound on performance, method out of sample likelihoods were compared to a distribution of likelihoods associated with a random phase inference method, where phases were drawn from a random circular uniform distribution. To build this distribution, *D*^(*m*)^ was computed 50 times for the random circular uniform approach. To contextualize how methods performed relative to this random approach, the difference between method log likelihoods and the median of the random log likelihood distribution were computed. These differences were then scaled by the standard deviation of the random log likelihood distribution, and are shown in Figures 3c and Supplementary Figures 15-17c.

For additional context, method likelihoods were also compared to likelihoods associated with treating the cell sample collection phases as the cell circadian phases, shown as the “LD” bar in Figures 3c and Supplementary Figures 15-17c.

### Selection of annotated cycling cell cycle genes

The 378 genes from CycleBase^17^ were considered as candidate core clock genes whose variation could describe cell cycle phase. This gene set was filtered in two ways. First, CycleBase genes were filtered for those with high variance. A mean-variance model was fit to all genes using counts from *Hsiao et al*., and CycleBase genes with a Pearson residual greater than 1 were kept. Second, CycleBase genes were filtered based on their correlation with two cell cycle markers: 1) Cdk1, a G2 marker 2) Cdc6, a G1/S phase marker. CycleBase genes with a Pearson correlation with Cdk1 greater than or equal to 0.2 and correlation with Cdc6 less than or equal to 0 were kept. In addition, CycleBase genes with a correlation with Cdc6 greater than or equal to 0.2 and correlation with Cdk1 less than or equal to 0 were kept. This yielded a set of 40 genes (Supplementary Table 3) treated as “core clock” cell cycle genes

### Hsiao et al. and Buettner et al. data cell cycle phase estimation

The acrophase priors used by Tempo for the core clock cell cycle genes were set as follows. Genes with CycleBase discrete peak time annotations of G1, G1/S, S, G2, G2/M, and M were assigned acrophase prior locations of 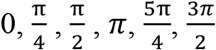 radians, respectively. The 95% interval width of each gene’s prior acrophase distribution was set to 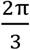 radians. Cdk1 was set as the reference gene.

Due to the asynchronous nature of the cell cycle, cell phases may not be symmetrically distributed over the circle for a given sample of cells. Under this setting, Tempo’s gene sinusoidal parameter estimates may not be reliable. Given this, Tempo was run without optimization of the core clock variational distributions in Step 1 of the algorithm. Instead, core clock gene variational distributions were fixed to the prior distributions (by setting the opt_phase_est_gene_params parameter in Tempo to False).

