## Supplemental Figures for "Tempo: an unsupervised Bayesian algorithm for circadian phase inference in single-cell transcriptomics"

**Supplementary Figures**

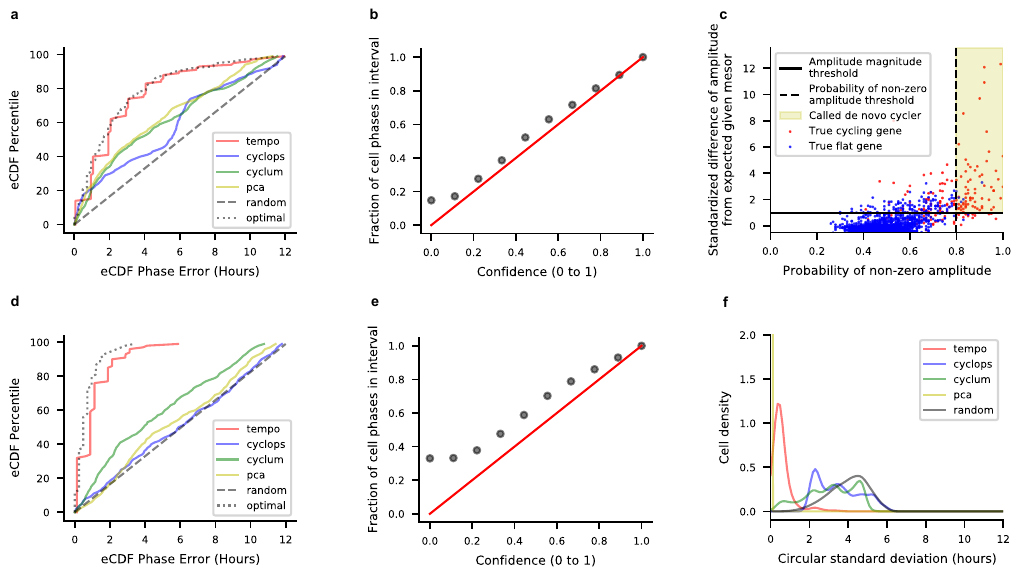

**Supplementary Figure 1:** Results on a simulated scRNA-seq dataset of 500 unsynchronized cells with mean library size of 10000 UMI. **a)** Empirical cumulative distribution function (eCDF) of the errors for each method’s cell phase point estimates, where all methods were run using the true core clock genes as input. **b)** Calibration of Tempo’s uncertainty estimates when run using the true core clock genes as input. **c)** Tempo’s *de novo* cycler detection procedure **d)** eCDF of the errors for method cell phase point estimates, where methods were run using default settings. **e)** Calibration of Tempo’s uncertainty estimates when run with default settings. **f)** Model stability when methods were run 5 times using default settings.

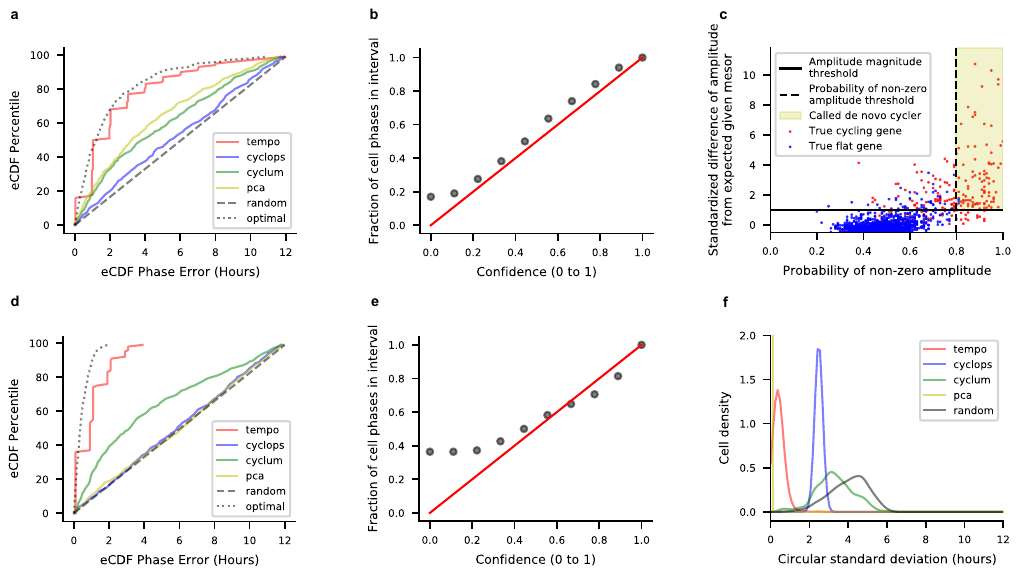

**Supplementary Figure 2:** Results on a simulated scRNA-seq dataset of 500 unsynchronized cells with mean library size of 20000 UMI. **a)** Empirical cumulative distribution function (eCDF) of the errors for each method’s cell phase point estimates, where all methods were run using the true core clock genes as input. **b)** Calibration of Tempo’s uncertainty estimates when run using the true core clock genes as input. **c)** Tempo’s *de novo* cycler detection procedure **d)** eCDF of the errors for method cell phase point estimates, where methods were run using default settings. **e)** Calibration of Tempo’s uncertainty estimates when run with default settings. **f)** Model stability when methods were run 5 times using default settings.

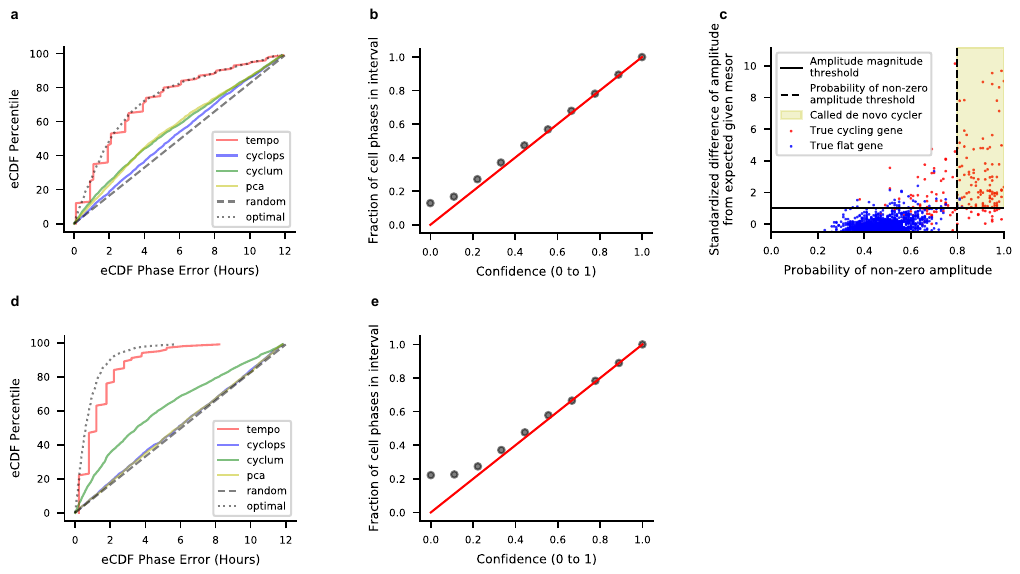

**Supplementary Figure 3:** Results on a simulated scRNA-seq dataset of 2000 unsynchronized cells with mean library size of 5000 UMI. **a)** Empirical cumulative distribution function (eCDF) of the errors for each method’s cell phase point estimates, where all methods were run using the true core clock genes as input. **b)** Calibration of Tempo’s uncertainty estimates when run using the true core clock genes as input. **c)** Tempo’s *de novo* cycler detection procedure **d)** eCDF of the errors for method cell phase point estimates, where methods were run using default settings. **e)** Calibration of Tempo’s uncertainty estimates when run with default settings.

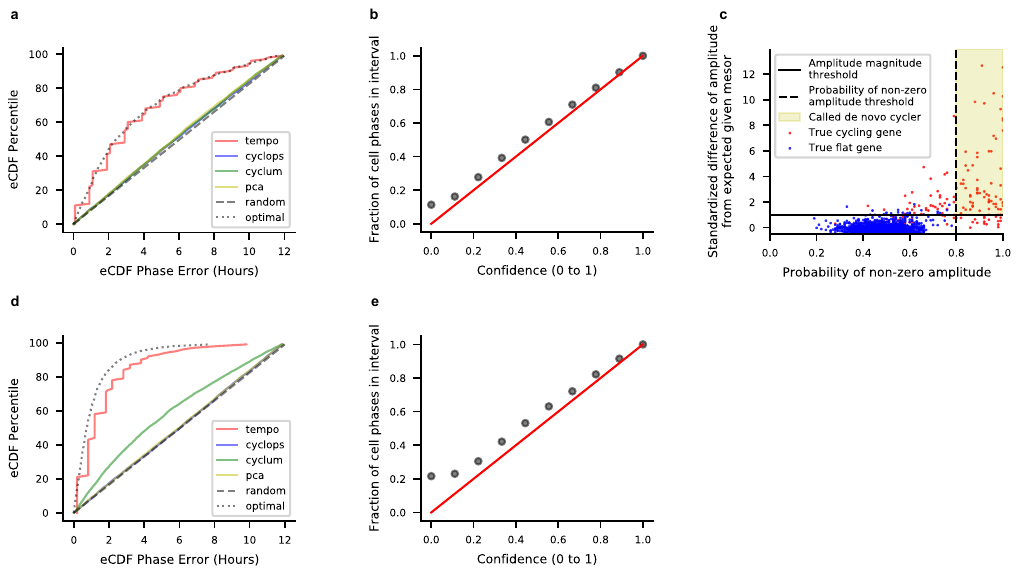

**Supplementary Figure 4:** Results on a simulated scRNA-seq dataset of 5000 unsynchronized cells with mean library size of 3000 UMI. **a)** Empirical cumulative distribution function (eCDF) of the errors for each method’s cell phase point estimates, where all methods were run using the true core clock genes as input. **b)** Calibration of Tempo’s uncertainty estimates when run using the true core clock genes as input. **c)** Tempo’s *de novo* cycler detection procedure **d)** eCDF of the errors for method cell phase point estimates, where methods were run using default settings. **e)** Calibration of Tempo’s uncertainty estimates when run with default settings.

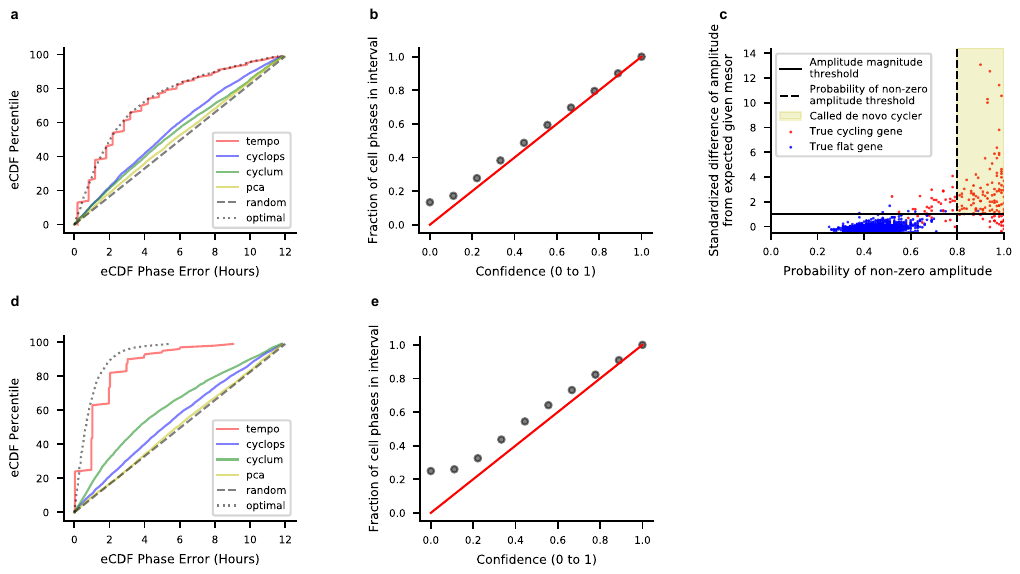

**Supplementary Figure 5:** Results on a simulated scRNA-seq dataset of 5000 unsynchronized cells with mean library size of 5000 UMI. **a)** Empirical cumulative distribution function (eCDF) of the errors for each method’s cell phase point estimates, where all methods were run using the true core clock genes as input. **b)** Calibration of Tempo’s uncertainty estimates when run using the true core clock genes as input. **c)** Tempo’s *de novo* cycler detection procedure **d)** eCDF of the errors for method cell phase point estimates, where methods were run using default settings. **e)** Calibration of Tempo’s uncertainty estimates when run with default settings.

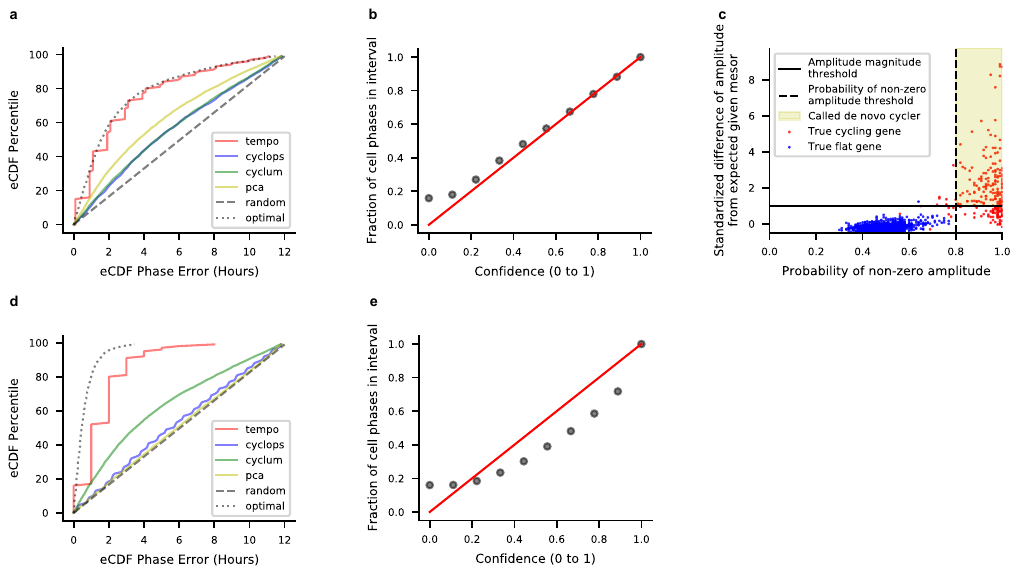

**Supplementary Figure 6:** Results on a simulated scRNA-seq dataset of 5000 unsynchronized cells with mean library size of 10000 UMI. **a)** Empirical cumulative distribution function (eCDF) of the errors for each method’s cell phase point estimates, where all methods were run using the true core clock genes as input. **b)** Calibration of Tempo’s uncertainty estimates when run using the true core clock genes as input. **c)** Tempo’s *de novo* cycler detection procedure **d)** eCDF of the errors for method cell phase point estimates, where methods were run using default settings. **e)** Calibration of Tempo’s uncertainty estimates when run with default settings.

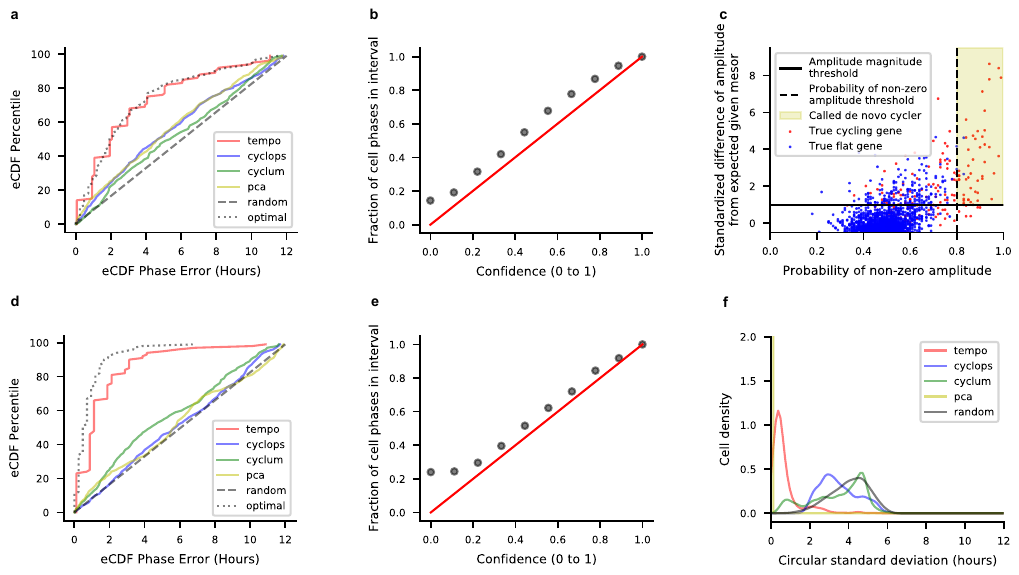

**Supplementary Figure 7:** Results on a simulated scRNA-seq dataset of 500 cells collected over a light-dark cycle (ZT0, ZT6, ZT12, ZT18) with mean library size of 5000 UMI. **a)** Empirical cumulative distribution function (eCDF) of the errors for each method’s cell phase point estimates, where all methods were run using the true core clock genes as input. **b)** Calibration of Tempo’s uncertainty estimates when run using the true core clock genes as input. **c)** Tempo’s *de novo* cycler detection procedure **d)** eCDF of the errors for method cell phase point estimates, where methods were run using default settings. **e)** Calibration of Tempo’s uncertainty estimates when run with default settings. **f)** Model stability when methods were run 5 times using default settings.

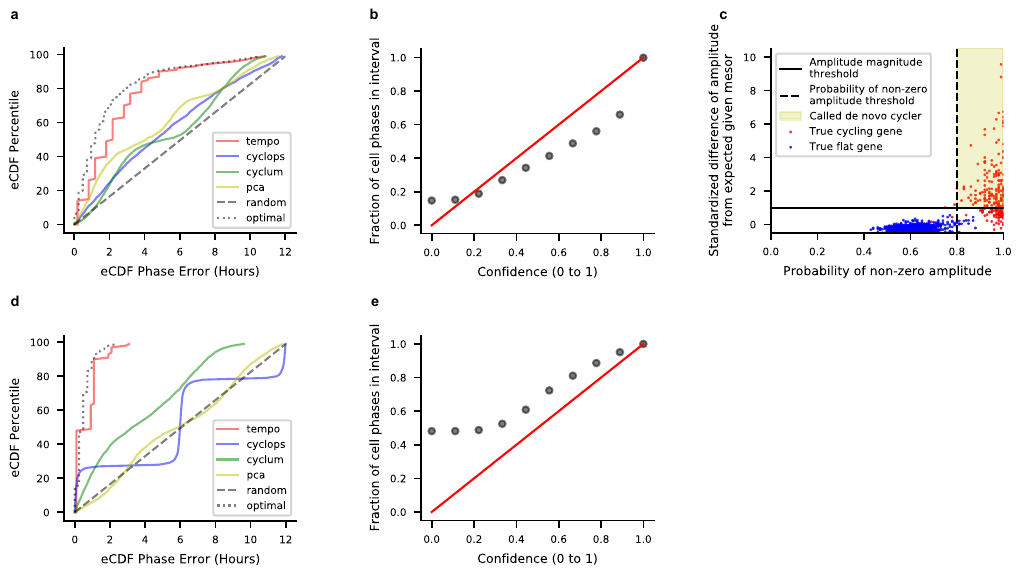

**Supplementary Figure 8:** Results on a simulated scRNA-seq dataset of 3000 cells collected over a light-dark cycle (ZT0, ZT6, ZT12, ZT18) with mean library size of 20000 UMI. **a)** Empirical cumulative distribution function (eCDF) of the errors for each method’s cell phase point estimates, where all methods were run using the true core clock genes as input. **b)** Calibration of Tempo’s uncertainty estimates when run using the true core clock genes as input. **c)** Tempo’s *de novo* cycler detection procedure **d)** eCDF of the errors for method cell phase point estimates, where methods were run using default settings. **e)** Calibration of Tempo’s uncertainty estimates when run with default settings.

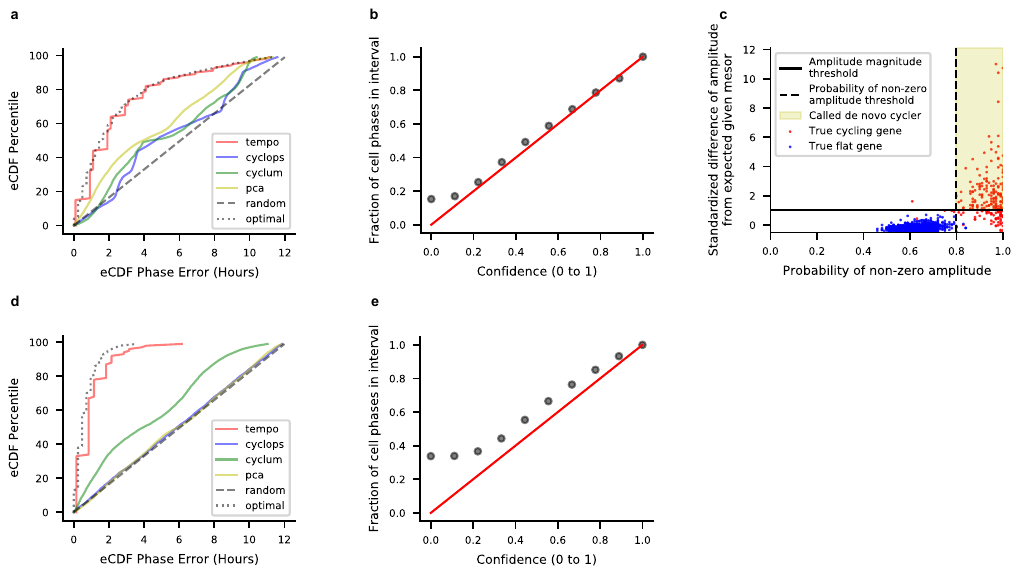

**Supplementary Figure 9:** Results on a simulated scRNA-seq dataset of 5000 cells collected over a light-dark cycle (ZT0, ZT6, ZT12, ZT18) with mean library size of 10000 UMI. **a)** Empirical cumulative distribution function (eCDF) of the errors for each method’s cell phase point estimates, where all methods were run using the true core clock genes as input. **b)** Calibration of Tempo’s uncertainty estimates when run using the true core clock genes as input. **c)** Tempo’s *de novo* cycler detection procedure **d)** eCDF of the errors for method cell phase point estimates, where methods were run using default settings. **e)** Calibration of Tempo’s uncertainty estimates when run with default settings.

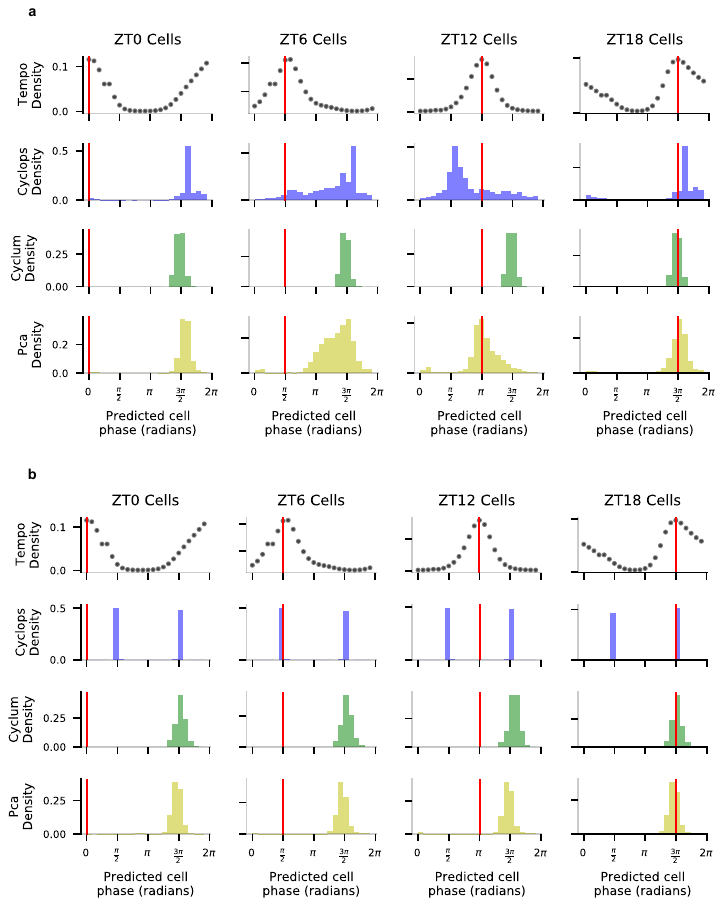

**Supplementary Figure 10**: Density of method cell phase predictions for aorta SMCs at various sample collection times. Tempo’s densities represent the pseudobulk approximate posterior distributions at each sample collection time point. Competing method densities represent method point estimates. a) Method cell phase predictions densities when run using only the core clock genes. b) Method cell phase predictions densities when run with default settings

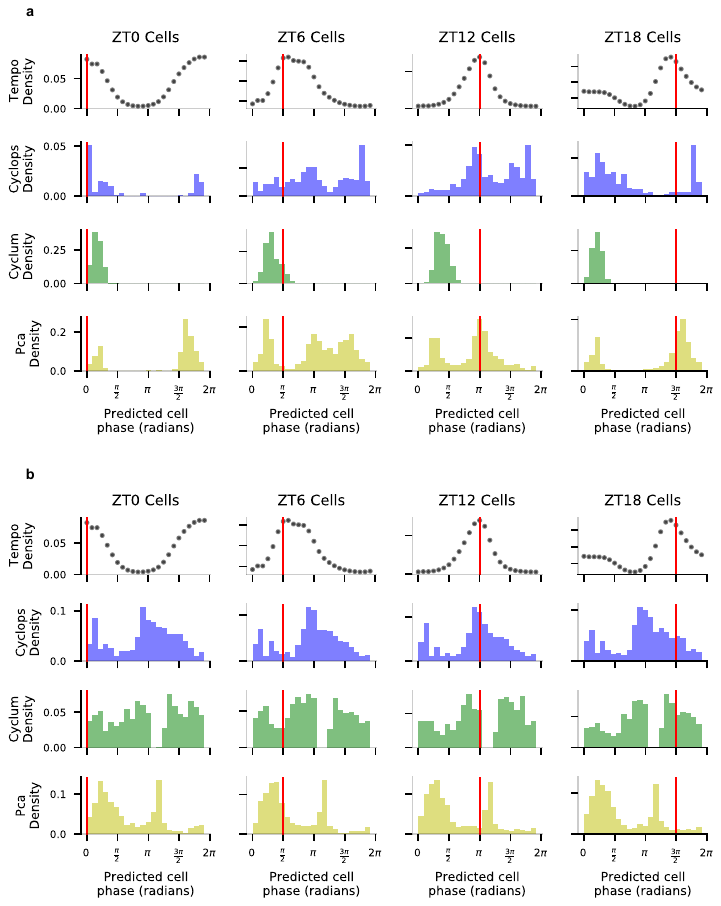

**Supplementary Figure 11**: Density of method cell phase predictions for aorta fibroblasts at various sample collection times. Tempo’s densities represent the pseudobulk approximate posterior distributions at each sample collection time point. Competing method densities represent method point estimates. a) Method cell phase predictions densities when run using only the core clock genes. b) Method cell phase predictions densities when run with default settings

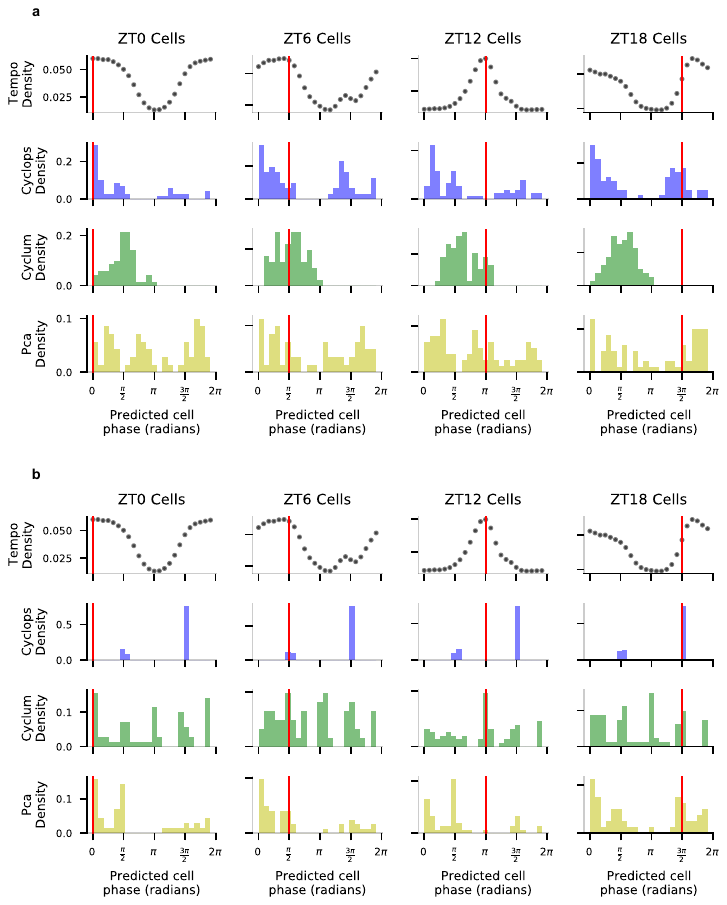

**Supplementary Figure 12**: Density of method cell phase predictions for aorta endothelial cells at various sample collection times. Tempo’s densities represent the pseudobulk approximate posterior distributions at each sample collection time point. Competing method densities represent method point estimates. a) Method cell phase predictions densities when run using only the core clock genes. b) Method cell phase predictions densities when run with default settings

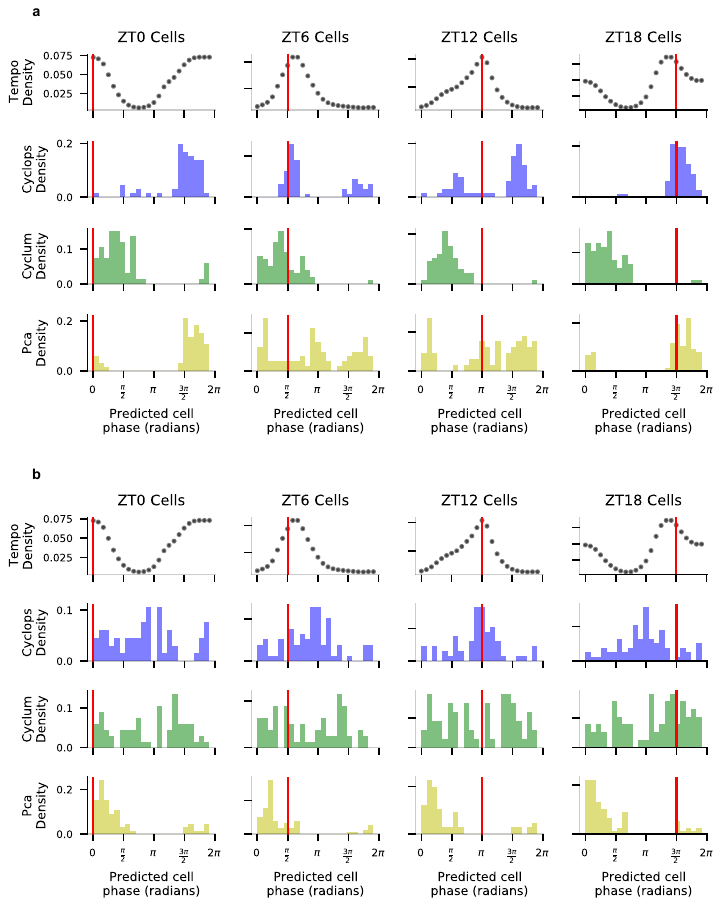

**Supplementary Figure 13**: Density of method cell phase predictions for aorta macrophages at various sample collection times. Tempo’s densities represent the pseudobulk approximate posterior distributions at each sample collection time point. Competing method densities represent method point estimates. a) Method cell phase predictions densities when run using only the core clock genes. b) Method cell phase predictions densities when run with default settings

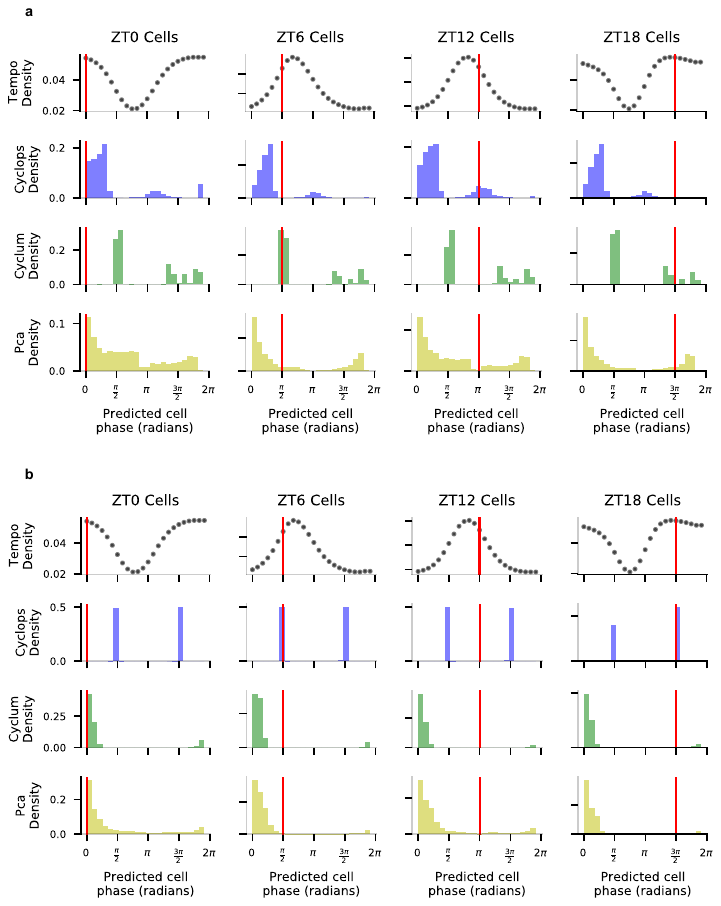

**Supplementary Figure 14**: Density of method cell phase predictions for liver hepatocytes at various sample collection times. Tempo’s densities represent the pseudobulk approximate posterior distributions at each sample collection time point. Competing method densities represent method point estimates. a) Method cell phase predictions densities when run using only the core clock genes. b) Method cell phase predictions densities when run with default settings

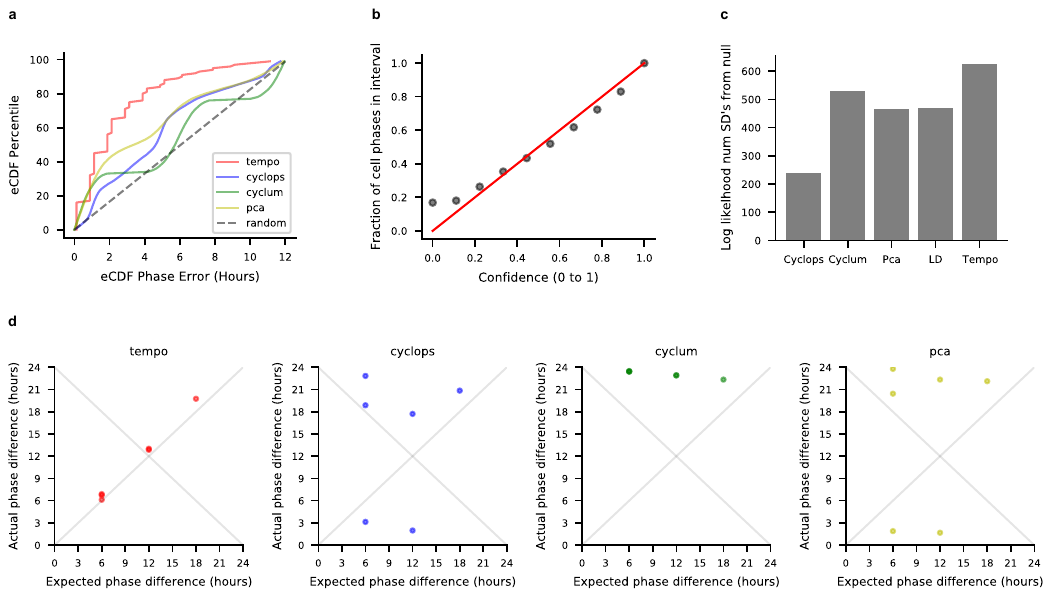

**Supplementary Figure 15:** Method results (core clock genes as input) on light-dark cycle aorta smooth muscle cells. Treating the sample collection phase in the light-dark cycle as the true cell circadian phases: **a)** eCDF of the errors for each method’s cell phase point estimates **b)** Calibration of Tempo’s uncertainty estimates. **c)** Method out of sample core clock gene likelihood analysis. LD corresponds to treating sample collection times as the true cell phases. Out of sample core clock likelihoods were computed for each method, and reported in terms of standard deviations from the median of a distribution of random likelihoods. **d)** Method relative shift analysis. Each dot represents a pair of sample collection times in the light-dark cycle (e.g. all 6 possible pairs of ZT0, ZT6, ZT12, ZT18), and conveys the relationship between the expected phase difference between a pair of time points and the actual phase difference for each method. As the phase difference is a circular random variable, methods with points lying along either y = x or y = 24 – x denote perfect performance.

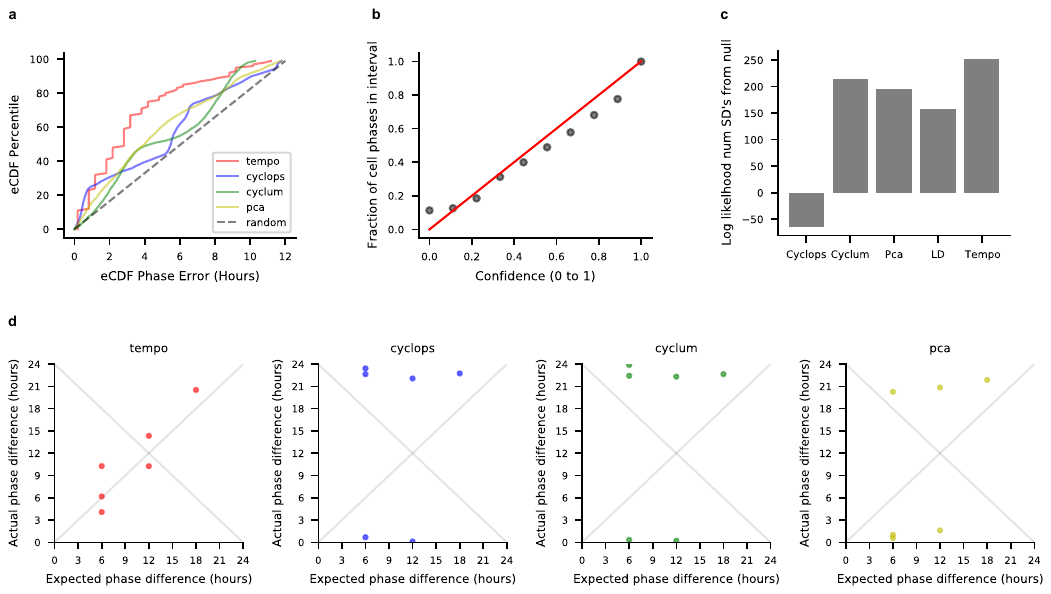

**Supplementary Figure 16:** Method results (clock genes as input) on light-dark cycle aorta fibroblasts. Treating the sample collection phase in the light-dark cycle as the true cell circadian phases: **a)** eCDF of the errors for each method’s cell phase point estimates **b)** Calibration of Tempo’s uncertainty estimates. **c)** Method out of sample core clock gene likelihood analysis. LD corresponds to treating sample collection times as the true cell phases.Out of sample core clock likelihoods were computed for each method, and reported in terms of standard deviations from the median of a distribution of random likelihoods. **d)** Method relative shift analysis. Each dot represents a pair of sample collection times in the light-dark cycle (e.g. all 6 possible pairs of ZT0, ZT6, ZT12, ZT18), and conveys the relationship between the expected phase difference between a pair of time points and the actual phase difference for each method. As the phase difference is a circular random variable, methods with points lying along either y = x or y = 24 – x denote perfect performance.

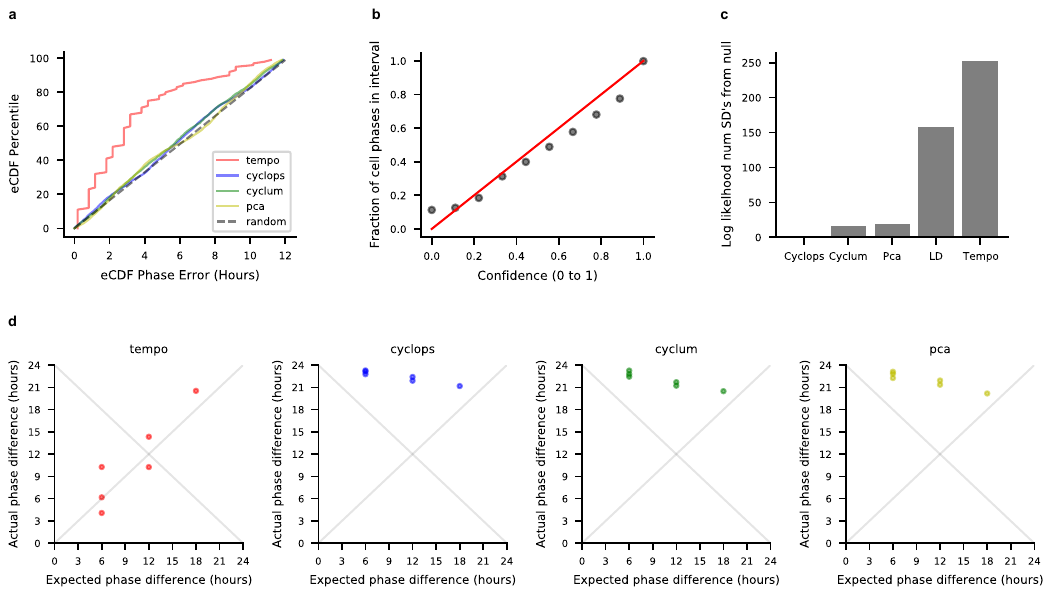

**Supplementary Figure 17:** Method results (default settings) on light-dark cycle aorta fibroblasts. Treating the sample collection phase in the light-dark cycle as the true cell circadian phases: **a)** eCDF of the errors for each method’s cell phase point estimates **b)** Calibration of Tempo’s uncertainty estimates. **c)** Method out of sample core clock gene likelihood analysis. LD corresponds to treating sample collection times as the true cell phases. Out of sample core clock likelihoods were computed for each method, and reported in terms of standard deviations from the median of a distribution of random likelihoods. **d)** Method relative shift analysis. Each dot represents a pair of sample collection times in the light-dark cycle (e.g. all 6 possible pairs of ZT0, ZT6, ZT12, ZT18), and conveys the relationship between the expected phase difference between a pair of time points and the actual phase difference for each method. As the phase difference is a circular random variable, methods with points lying along either y = x or y = 24 – x denote perfect performance.

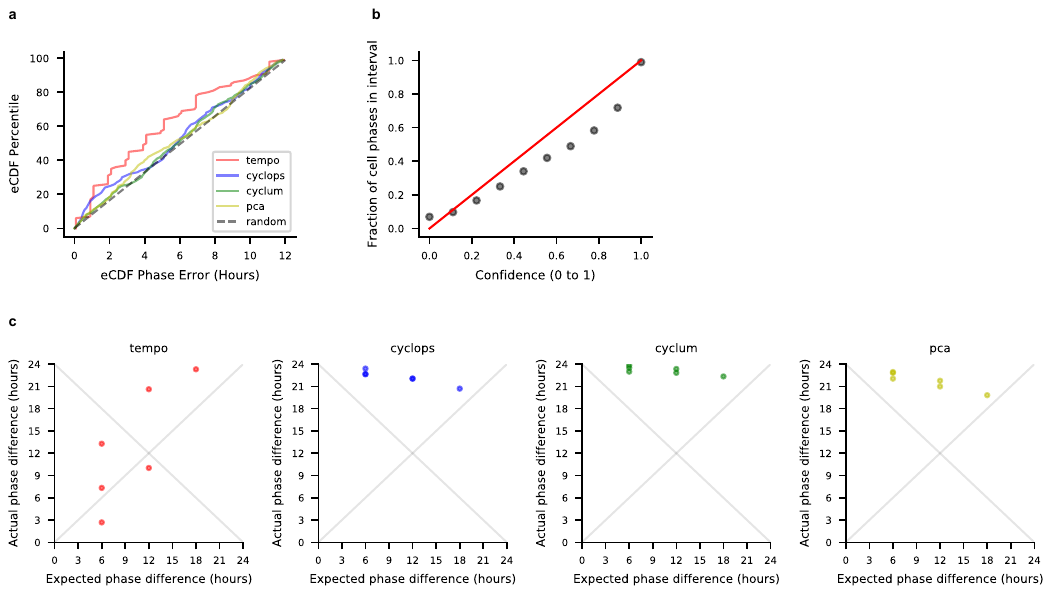

**Supplementary Figure 18:** Method results (clock genes as input) on light-dark cycle aorta endothelial cells. Treating the sample collection phase in the light-dark cycle as the true cell circadian phases: **a)** eCDF of the errors for each method’s cell phase point estimates **b)** Calibration of Tempo’s uncertainty estimates. **c)** Method relative shift analysis. Each dot represents a pair of sample collection times in the light-dark cycle (e.g. all 6 possible pairs of ZT0, ZT6, ZT12, ZT18), and conveys the relationship between the expected phase difference between a pair of time points and the actual phase difference for each method. As the phase difference is a circular random variable, methods with points lying along either y = x or y = 24 – x denote perfect performance.

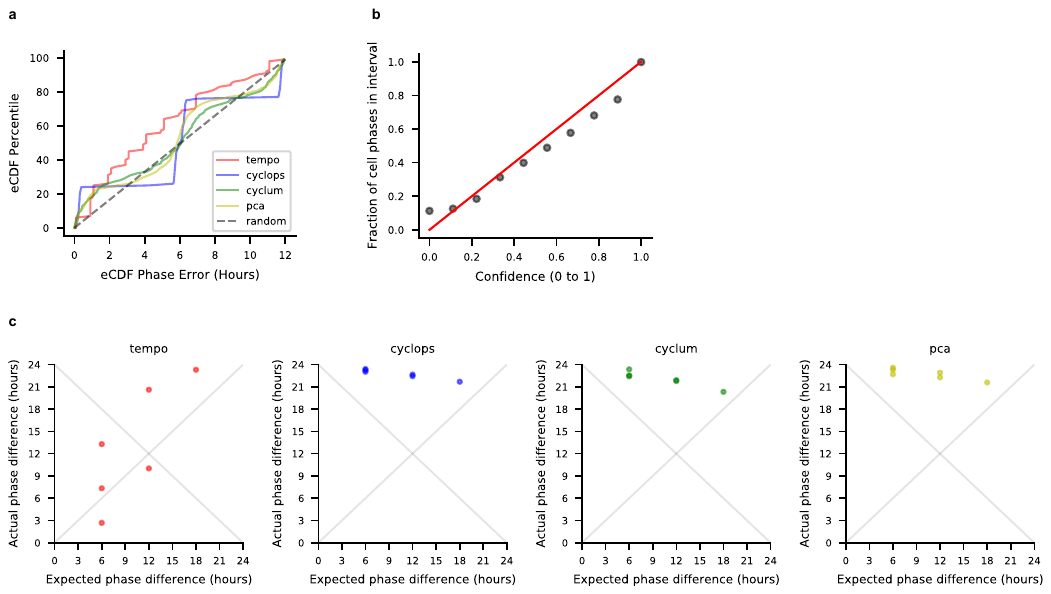

**Supplementary Figure 19:** Method results (default settings) on light-dark cycle aorta endothelial cells. Treating the sample collection phase in the light-dark cycle as the true cell circadian phases: **a)** eCDF of the errors for each method’s cell phase point estimates **b)** Calibration of Tempo’s uncertainty estimates. **c)** Method relative shift analysis. Each dot represents a pair of sample collection times in the light-dark cycle (e.g. all 6 possible pairs of ZT0, ZT6, ZT12, ZT18), and conveys the relationship between the expected phase difference between a pair of time points and the actual phase difference for each method. As the phase difference is a circular random variable, methods with points lying along either y = x or y = 24 – x denote perfect performance.

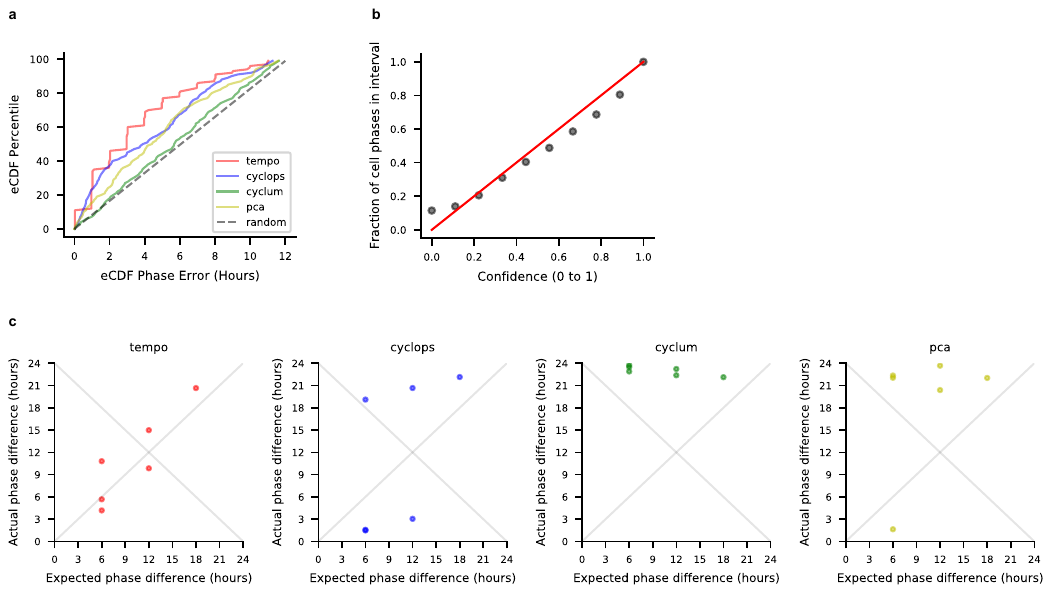

**Supplementary Figure 20:** Method results (clock genes as input) on light-dark cycle aorta macrophages. Treating the sample collection phase in the light-dark cycle as the true cell circadian phases: **a)** eCDF of the errors for each method’s cell phase point estimates **b)** Calibration of Tempo’s uncertainty estimates. **c)** Method relative shift analysis. Each dot represents a pair of sample collection times in the light-dark cycle (e.g. all 6 possible pairs of ZT0, ZT6, ZT12, ZT18), and conveys the relationship between the expected phase difference between a pair of time points and the actual phase difference for each method. As the phase difference is a circular random variable, methods with points lying along either y = x or y = 24 – x denote perfect performance.

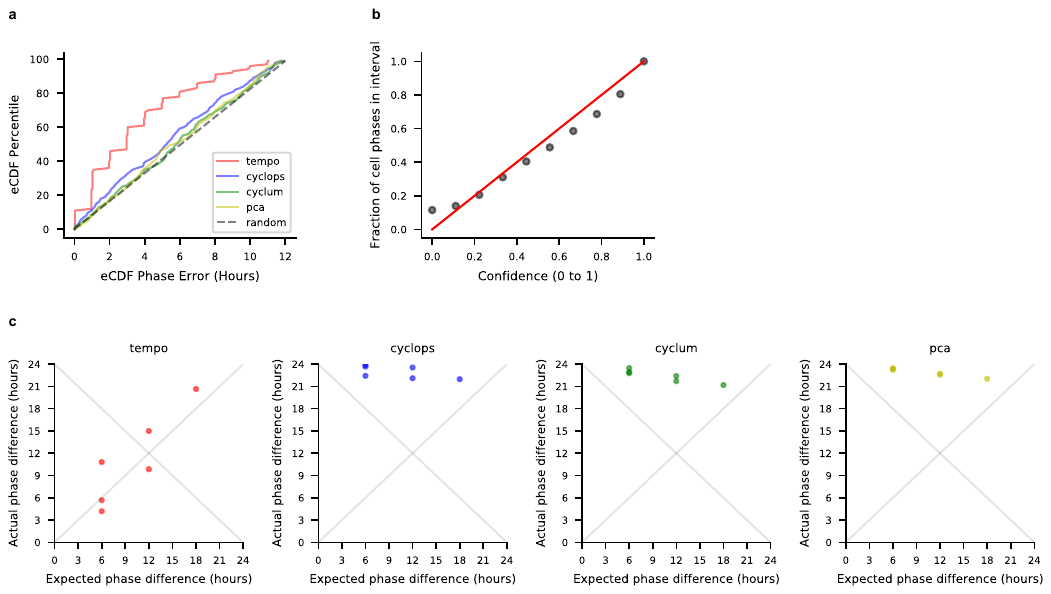

**Supplementary Figure 21:** Method results (default settings) on light-dark cycle aorta macrophages. Treating the sample collection phase in the light-dark cycle as the true cell circadian phases: **a)** eCDF of the errors for each method’s cell phase point estimates **b)** Calibration of Tempo’s uncertainty estimates. **c)** Method relative shift analysis. Each dot represents a pair of sample collection times in the light-dark cycle (e.g. all 6 possible pairs of ZT0, ZT6, ZT12, ZT18), and conveys the relationship between the expected phase difference between a pair of time points and the actual phase difference for each method. As the phase difference is a circular random variable, methods with points lying along either y = x or y = 24 – x denote perfect performance.

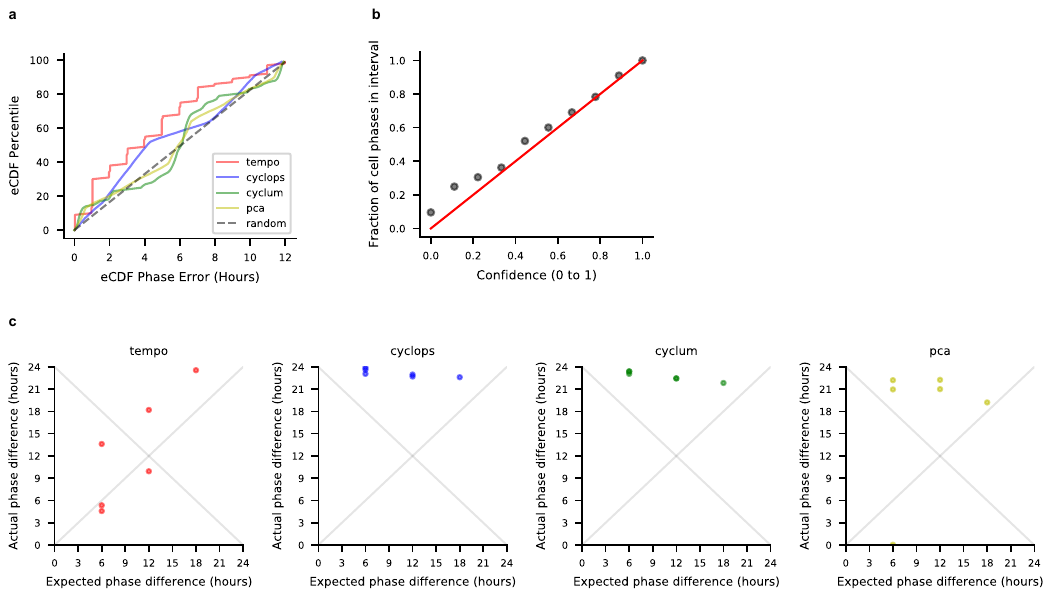

**Supplementary Figure 22:** Method results (clock genes as input) on light-dark cycle liver hepatocytes. Treating the sample collection phase in the light-dark cycle as the true cell circadian phases: **a)** eCDF of the errors for each method’s cell phase point estimates **b)** Calibration of Tempo’s uncertainty estimates. **c)** Method relative shift analysis. Each dot represents a pair of sample collection times in the light-dark cycle (e.g. all 6 possible pairs of ZT0, ZT6, ZT12, ZT18), and conveys the relationship between the expected phase difference between a pair of time points and the actual phase difference for each method. As the phase difference is a circular random variable, methods with points lying along either y = x or y = 24 – x denote perfect performance.

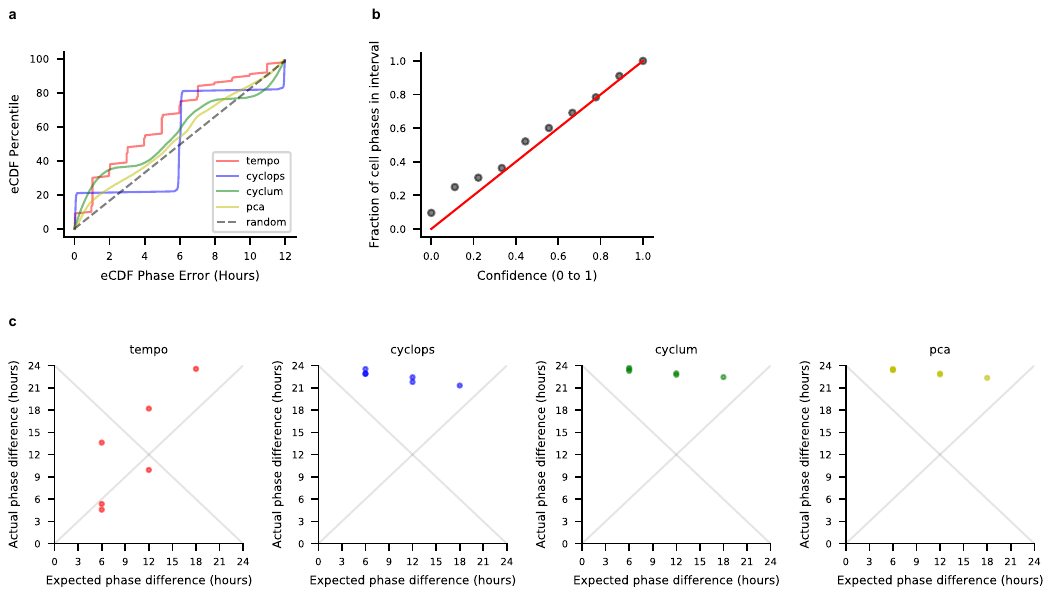

**Supplementary Figure 23:** Method results (default settings) on light-dark cycle liver hepatocytes. Treating the sample collection phase in the light-dark cycle as the true cell circadian phases: **a)** eCDF of the errors for each method’s cell phase point estimates **b)** Calibration of Tempo’s uncertainty estimates. **c)** Method relative shift analysis. Each dot represents a pair of sample collection times in the light-dark cycle (e.g. all 6 possible pairs of ZT0, ZT6, ZT12, ZT18), and conveys the relationship between the expected phase difference between a pair of time points and the actual phase difference for each method. As the phase difference is a circular random variable, methods with points lying along either y = x or y = 24 – x denote perfect performance.
