## Supplemental Information and Methods for "Tempo: an unsupervised Bayesian algorithm for circadian phase inference in single-cell transcriptomics"

### Tempo Supplementary Information

#### 1 Estimating the transcript proportion - dispersion relationship

Tempo uses a Negative Binomial (NB) data likelihood model, as it can account for the overdispersion observed in real count data. While many existing applications of NB to scRNA-seq data model dispersions in a gene-specific fashion, these estimates are known to exhibit high amounts of uncertainty. Methods such as scTransform and DESeq2 combat this by an empirical bayes approach to shrink dispersion estimates, assuming genes with similar means share similar dispersions. We take this a step further and make the simplifying assumption that the transcript proportion - dispersion relationship across all genes is strictly described by a polynomial function $g_\zeta(\lambda_{ij})$ parameterized by $\zeta$. The coefficients of this polynomial, $\zeta$, are estimated as follows:

##### Step 1: Estimate gene-specific proportions and dispersions

We presume gene counts are distributed according to:

$$X_{ij} \sim NB(\tilde{\lambda_j} * L_i, \tilde{\delta_j}) \tag{1}$$

Where:

$$E[X_{ij}] = \tilde{\lambda_j} * L_i \tag{2}$$

$$Var(X_{ij}) = E[X_{ij}] + \tilde{\delta_j}(E[X_{ij}])^2 \tag{3}$$

and $\tilde{\lambda_j}$ and $\tilde{\delta_j}$ are the temporary (i.e. only used to estimate $\zeta$) gene-specific transcript proportion and dispersion for gene $j$, respectively. Under this likelihood model, for each gene we estimate the maximum likelihood transcript proportion $\tilde{\lambda_j}$ and dispersion $\tilde{\delta_j}$.

##### Step 2: Fit polynomial model describing the transcript proportion - dispersion relationship across all genes

Assume g is a polynomial function whose coefficients $\zeta$ defines the transcript log proportion - log dispersion relationship shared across all genes:

$$g_\zeta(\lambda_{ij}) = log(\delta_j) = \sum_{k=1}^{||\zeta||} \zeta_k log^k(\lambda_{ij}) \tag{4}$$

Using OLS, we estimate the coefficients $\zeta$ under the following objective:

$$\arg\min_{\zeta} \sum_{j=1}^{p} (log(\tilde{\delta_j}) - g_\zeta(\tilde{\lambda_j}))^2 \tag{5}$$

where $p$ is the number of genes, and $\tilde{\lambda_j}$ and $\tilde{\delta_j}$ are the temporary gene-specific transcript proportion and dispersion for gene $j$ estimated in Step 1.

##### Note

We note that dispersion estimates may be upwardly biased as a subset of genes cycling over the circadian cycle will be fit under a mean model that assumes flat expression over the circadian cycle. However, we assume the fraction of cycling genes detected is small, and thus should not upwardly bias the dispersion estimates much.

#### 2 Tempo approximate generative model

We model $P(\theta, \beta|X)$ as an approximate distribution $q(\theta, \beta)$. In practice, we factor the joint $q(\theta, \beta)$ into a conditional and a marginal:

$$q(\theta, \beta) = q(\theta|\beta)q(\beta) \tag{6}$$

#### 2.1 $q(\beta)$

$q(\beta)$ is the approximate marginal posterior with respect to the gene parameters. $q(\beta)$ is parameterized by $\tilde{\beta} = (\tilde{\beta}_1, ..., \tilde{\beta}_p)$. $\tilde{\beta}_j$ refers to the parameters for gene $j$ and $\tilde{\beta}_j = (\tilde{\mu_j}^{(loc)}, \tilde{\mu_j}^{(scale)}, \tilde{A_j}^{(\alpha)}, \tilde{A_j}^{(\beta)}, \tilde{\phi_j}^{(loc)}, \tilde{\phi_j}^{(scale)}, \tilde{\gamma_j}^{(\alpha)}, \tilde{\gamma_j}^{(\beta)}, \tilde{C_j})$. Approximate samples $\beta_j^* = (\mu_j^*, A_j^*, \phi_j^*, Q_j^*)$ are drawn according to the following generative process:

$$\mu_j^* \sim Normal(\tilde{\mu_j}^{(loc)}, \tilde{\mu_j}^{(scale)}) \tag{7}$$

$$A_j^* \sim \tilde{C_j} * Beta(\tilde{A_j}^{(\alpha)}, \tilde{A_j}^{(\beta)}) * (A^{(max)} - A^{(min)}) + A^{(min)} \tag{8}$$

$$\phi_j^* \sim PowerSpherical(\tilde{\phi_j}^{(loc)}, \tilde{\phi_j}^{(scale)}) \tag{9}$$

$$\gamma_j^* \sim Beta(\tilde{\gamma_j}^{(\alpha)}, \tilde{\gamma_j}^{(\beta)}) \tag{10}$$

$$Q_j^* \sim Bernoulli(\gamma_j^*). \tag{11}$$

$A^{(min)}$ and $A^{(max)}$ are the same as the values used in specifying the priors for A. $\tilde{C_j}$ is an indicator variable, equal to 1 if a gene is current cycling gene (i.e. core clock gene or *de novo* cycling gene). This allows Tempo's approximate generative model to model non-cycling genes as having flat expression over the circadian cycle. Of note, we use a Power Spherical distribution (*De Cao et al.*, see citation 32 in manuscript) to model the approximate posterior of $\phi$, as it has similar shape to the Von Mises distribution but has a differentiable implementation in PyTorch that is amenable to optimization of its parameters.

#### 2.2 $q(\theta|\beta)$

$P(\theta|X, \beta)$ can be well approximated using grid sampling. As such, Tempo models $q(\theta|\beta)$ as an approximation of $P(\theta|X, \beta)$ using grid sampling. Dividing the domain of $\theta$ into k equidistant points on [0,2$\pi$]:

$$q(\theta_i = \frac{2\pi}{k}\nu|\beta) = \frac{P(X_i|\theta_i = \frac{2\pi}{k}\nu, \beta)P(\theta_i)}{\sum_{\nu^*=0}^{k-1} P(X_i|\theta_i = \frac{2\pi}{k}\nu^*, \beta)P(\theta_i)} \tag{12}$$

Of note, as $k$ approaches $\infty$, $q(\theta|\beta)$ converges to $P(\theta|X, \beta)$. Moreover, in practice $q(\theta|\beta)$ is computed using approximate samples of $\beta$ from $q(\beta)$. Given this, only genes for which $\tilde{C_j} = 1$ contribute information to the estimation of $q(\theta|\beta)$.

### 3 Tempo algorithm to minimize the KL between $q(\theta, \beta)$ and $P(\theta, \beta|X)$

#### 3.1 Initialization of $q(\theta, \beta)$

Prior to the start of the algorithm, Tempo filters genes to reduce the computational burden of *de novo* cycler detection. First, genes are filtered by their pseudobulk proportion according to a user-specified threshold ($10^{-5}$ by default). Second, genes are filtered for having low variance. *A priori*, such genes are unlikely to be cycling genes. A mean-variance model is fit to non-clock genes and then genes with small Pearson residuals are discarded. For more detail on this filtering of candidate *de novo* cyclers, please see section 4.

Using the remaining genes, $\tilde{\beta}$ is initialized to minimize the KL divergence between $q(\beta)$ and $P(\beta)$. Moreover, cycling gene indicators are set such that $\tilde{C_j} = 1$ for known core clock genes, and $\tilde{C_j} = 0$ for non-core clock genes.

#### 3.2 Step 1: estimation of cell phase distribution using current cycling genes

Tempo minimizes the KL divergence between $q(\theta, \beta)$ and $P(\theta, \beta|X)$ through minimizing the following evidence lower bound (ELBO) loss function (the derivation of which can be viewed in section 5 below):

$$ELBO(\tilde{\beta}) = KL(q(\beta)||P(\beta)) - E_{q(\theta,\beta)}[logP(X|\theta, \beta)] \tag{13}$$

where $\tilde{\beta}$ denotes the parameters describing the shape of $q(\beta)$. $\tilde{\beta}$ is optimized using stochastic gradient descent. Of note $q(\theta|\beta)$ is completely determined by $q(\beta)$ and X for our model, as current cycling genes only contribute to the estimate of $\theta$ in $q(\theta|\beta)$. Moreover, when computing the objective function that is minimized in equation 14, the expectations are computed using a Monte-Carlo approximation.

##### 3.3 Step 2: identification of *de novo* cycling genes

Optimal estimation of cell phase should make use of information from both the core clock genes and cell-type-specific CCGs. However, CCGs are often unknown ahead of time. In Step 2, we use a heuristic procedure to identify *de novo* cycling genes with expression that are well-explained by the current approximate cell phase distribution $q(\theta|\beta)$. To do this, we consider the current genes $j$ for which $\tilde{C}_j = 0$. For this gene set, approximate gene parameter distributions are fit by optimizing equation 14 assuming the genes are cyclers (by fixing $\tilde{C}_j = 1$) and conditional on the cell phase distributions computed in Step 1.

Based on the approximate posteriors for $\gamma$ and A, genes with high posterior probabilities of having non-zero amplitude and genes with sufficiently high amplitude are called *de novo* cyclers, and $\tilde{C}_j$ is set to 1 for such genes. For genes not satisfying these criteria, $\tilde{C}_j = 0$. More details on the criteria used to call *de novo* cycling genes from $\tilde{\mu}$, $\tilde{A}$, $\tilde{\phi}$, and $\tilde{\gamma}$ can be found in section 8.

##### 3.4 Algorithm progression

Using information from the core clock genes alone, Tempo uses Step 1 to approximate the latent cell phase distribution. Tempo then tests whether the latent cell phase distribution explains the data better than random and halts the algorithm progression if it does not (details of which can be viewed in sections 6 and 7). Using the latent cell phase distribution, in Step 2, Tempo identifies *de novo* cycling genes. Tempo then re-estimates the latent cell phase distribution using information from both the core clock and *de novo* cycling genes. At each iteration, Tempo alternates between Step 1 and Step 2 until the evidence for the core clock expression worsens (computation of which can be viewed in section 6) or the algorithm exceeds the maximum of iterations specified by the user. Tempo returns $\tilde{\beta}$, and corresponding $q(\beta)$ and $q(\theta|\beta)$, from the last iteration for which the evidence for the core clock expression is at least as good as the evidence from the first iteration.

#### 4 Optional preprocessing step: restrict the data to highly variable genes

To improve computational efficiency, Tempo offers (and recommends) the option to restrict the data to highly variable genes with outlier variances, as *a priori* these genes may be the most likely to cycle over the circadian cycle and provide the most information about cell phase. First, a transformation, Z, of the count matrix, X, is computed as follows:

$$Z_{ij} = log_{10}(\frac{X_{ij} + 1}{L_i}) \tag{14}$$

Gene means and variances are then computed using the transformed matrix, Z. A univariate gaussian kernel (bandwidth 0.1, by default) is then used to learn the relationship between the transformed means and variances. Highly variable genes are then identified as those with a pearson residual greater than a user specified threshold, which is set to 0.5 standard deviations, by default.

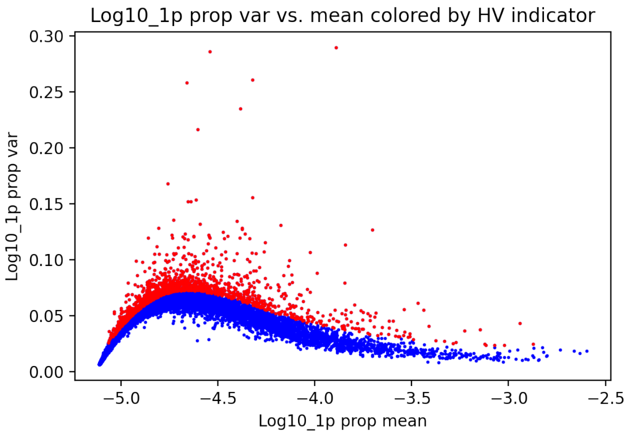


Figure 1: Example of the mean-variance relationship on the transformed data, where genes identified as highly variable are colored in red.

### 5 Objective function derivation

$$KL(q(\theta,\beta)||P(\theta,\beta|X)) \tag{15}$$

$$= \int_\beta \int_\theta q(\theta,\beta)log(\frac{q(\theta,\beta)}{P(\theta,\beta|X)}) \tag{16}$$

$$= \int_\beta \int_\theta q(\beta)q(\theta|\beta)log(\frac{q(\theta|\beta)q(\beta)}{P(\theta,\beta|X)}) \tag{17}$$

$$= \int_\beta \int_\theta q(\beta)q(\theta|\beta)log(\frac{q(\theta|\beta)q(\beta)}{P(\theta,\beta|X)} * \frac{\int_{\theta'} P(\theta',\beta|X)}{\int_{\theta'} P(\theta',\beta|X)}) \tag{18}$$

$$= \int_\beta q(\beta) \int_\theta q(\theta|\beta)log(\frac{q(\theta|\beta)\int_{\theta'} P(\theta',\beta|X)}{P(\theta,\beta|X)}) + \int_\beta q(\beta) \int_\theta q(\theta|\beta)log(\frac{q(\beta)}{\int_{\theta'} P(\theta',\beta|X)}) \tag{19}$$

We note that $\int_{\theta'} P(\theta',\beta|X)$ is that posterior marginal of $\beta$ and that $\frac{P(\theta,\beta|X)}{\int_{\theta'} P(\theta',\beta|X)}$ is the conditional posterior distribution of $\theta$ given $\beta$.

It can be show that:

$$\frac{P(\theta,\beta|X)}{\int_{\theta'} P(\theta',\beta|X)} = \frac{P(X|\theta,\beta)P(\theta)}{\int_{\theta'} P(X|\theta',\beta)P(\theta')} \tag{20}$$

Given this, the left term of equation (20) can be rewritten:

$$= \int_\beta q(\beta) \int_\theta q(\theta|\beta)log(\frac{q(\theta|\beta)\int_{\theta'} P(X|\theta',\beta)P(\theta')}{P(X|\theta,\beta)P(\theta)}) + \int_\beta q(\beta) \int_\theta q(\theta|\beta)log(\frac{q(\beta)}{\int_{\theta'} P(\theta',\beta|X)}) \tag{21}$$

We note that the left term is the KL divergence between $q(\theta|\beta)$ and the distribution (the conditional posterior distribution of $\theta$ given $\beta$) that it aims to approximate using grid sampling. As such, the KL divergence between these terms will go to zero (i.e. the approximation will be exact) as the number of grid points goes to $\infty$. Given that the definition of our model for $q(\theta|\beta)$ inherently minimizes the KL divergence of the left term, we only need to optimize parameters of $q(\beta)$ that minimize the right term.

Continuing to simplify the right term in equation (22):

$$\int_\beta q(\beta) \int_\theta q(\theta|\beta)log(\frac{q(\beta)}{\int_{\theta'} P(\theta',\beta|X)}) \tag{22}$$

$$= \int_\beta q(\beta) \int_\theta q(\theta|\beta)log(\frac{q(\beta)P(X)}{P(\beta)\int_{\theta'} P(X|\theta',\beta)P(\theta')}) \tag{23}$$

Pulling P(X) out of the log:

$$= log(P(X)) + \int_\beta q(\beta) \int_\theta q(\theta|\beta)log(\frac{q(\beta)}{P(\beta)\int_{\theta'} P(X|\theta',\beta)P(\theta')}) \tag{24}$$

Pulling $q(\beta)$ and $P(\beta)$ out of the log:

$$= log(P(X)) + KL(q(\beta)||P(\beta)) - \int_\beta q(\beta) \int_\theta q(\theta|\beta)log(\int_{\theta'} P(X|\theta',\beta)P(\theta')) \tag{25}$$

Using Jensen's inequality:

$$\leq log(P(X)) + KL(q(\beta)||P(\beta)) - \int_\beta q(\beta) \int_\theta q(\theta|\beta) \int_{\theta'} P(\theta')log(P(X|\theta',\beta)) \tag{26}$$

We note $\int_{\theta'} P(\theta')log(P(X|\theta',\beta))$ is the expected log probability of the data where $\theta$ comes from the prior distribution. Of note, $q(\theta|\beta)$ is defined as the grid approximation of the posterior of $\theta$ conditional on $\beta$, whose density values are determined by both the prior for $\theta$ and the probability of the data. As such, $\int_{\theta'} P(\theta')log(P(X|\theta',\beta)) \leq \int_{\theta'} q(\theta'|\beta)log(P(X|\theta',\beta))$. As such:

$$\begin{aligned} &log(P(X)) + KL(q(\beta)||P(\beta)) - \int_\beta q(\beta) \int_\theta q(\theta|\beta) \int_{\theta'} P(\theta')log(P(X|\theta',\beta)) \\ &\geq log(P(X)) + KL(q(\beta)||P(\beta)) - \int_\beta q(\beta) \int_\theta q(\theta|\beta) \int_{\theta'} q(\theta'|\beta)log(P(X|\theta',\beta)) \end{aligned} \tag{27}$$

Which simplifies to:

$$= log(P(X)) + KL(q(\beta)||P(\beta)) - \int_{\beta} q(\beta) \int_{\theta} q(\theta|\beta) log(P(X|\theta, \beta)) \tag{28}$$

$$= log(P(X)) + KL(q(\beta)||P(\beta)) - E_{q(\theta,\beta)}[log(P(X|\theta, \beta))] \tag{29}$$

$P(X)$ does not depend on the approximate distribution parameters. Therefore, we can ignore it when computing our ultimate objective function, ELBO($\tilde{\beta}$):

$$ELBO(\tilde{\beta}) = KL(q(\beta)||P(\beta)) - E_{q(\theta,\beta)}[logP(X|\theta, \beta)] \tag{30}$$

Where $\tilde{\beta}$ are the differentiable parameters describing the shape of $q(\beta)$). Minimizing this objective function will minimize the KL divergence between our approximate joint posterior and the true joint posterior.

### 6 Core clock Bayesian evidence computation

For a given $\tilde{\beta}$ and corresponding $q(\theta, \beta)$, core clock Bayesian evidence is computed as:

$$\int P(X^{(cc)}|\beta, \theta) q(\theta, \beta) \tag{31}$$

Where $X^{(cc)}$ denotes the cell transcript count matrix for the core clock genes. In practice, the Bayesian evidence is computed using a Monte-Carlo estimate.

### 7 Assessing whether $q(\theta, \beta)$ explains core clock expression better than random

After optimization of $\tilde{\beta}$ in Step 1, Tempo assesses whether the current approximate cell phase distributions explain core clock expression better than random. To do this, Tempo generates a random UMI matrix, $X^{(random)}$, where gene transcript counts are independently permuted across cells. Tempo then runs Step 1 on these data to fit $\tilde{\beta}^{(random)}$. The core clock Bayesian evidence (computation of which is described in section 6) associated with $\tilde{\beta}$ is then compared to that of $\tilde{\beta}^{(random)}$ via a Bayes factor. If the Bayes factor does not exceed a user-defined threshold (1.5 by default), this suggests not enough information exists (for technical or biological reasons) to sufficiently estimate cell phase and halts the algorithm progression.

### 8 *De novo* cycler selection criteria

Upon fitting $\tilde{\beta}$ in Step 2, Tempo identifies *de novo* cycling genes according to two criteria. First, Tempo selects genes better explained by a sinusoidal mean model than a flat mean model based on MAP values for Q. By default, genes are required to have MAP Q values of 0.8 or greater to be considered a *de novo* cycler. Second, Tempo selects genes with high amplitude conditional on the mesor. For current non-cyclers $j$, Tempo fits a Nadaraya-Watson kernel regression model $f$ describing the relationship between $\mu_j^{(MAP)}$ and $A_j^{(MAP)}$:

$$f(\mu_j^{(MAP)}) = \hat{A_j}^{(MAP)} \tag{32}$$

where $\hat{A_j}^{(MAP)}$ is the expected MAP amplitude of a gene given its MAP mesor. Tempo then computes the pearson residual of each gene's MAP amplitude, $t_j$, as follows:

$$t_j = \frac{A_j^{(MAP)} - \hat{A_j}^{(MAP)}}{\sigma} \tag{33}$$

Where:

$$\sigma = \sqrt{\frac{1}{p} \sum_{j=1}^{p} (A_j^{(MAP)} - \hat{A_j}^{(MAP)})^2} \tag{34}$$

and $p$ is the number of non-cyclers. By default, Tempo requires $t_j$ greater than 1 for genes to be considered a *de novo* cycler.

### 9 Data preprocessing and run settings for competing methods

*Cyclops*
For a given input transcript count matrix, genes with transcript pseudobulk proportions of $10^{-5}$ were kept. Transcript counts were then transformed by adding a pseudocount of 1 transcript, library size normalized, logged, and z-scored. The transformed count matrix was then transformed using PCA using as many principal components needed to explain at least 97% of the data variance. Principal components were then z-scored, and used as input to the Cyclum python package implementation of Cyclops.

*Cyclum*
For a given input transcript count matrix, genes with transcript pseudobulk proportions of $10^{-5}$ were kept. Transcript counts were then transformed by adding a pseudocount of 1 transcript, library size normalized, logged, and z-scored. The transformed count matrix was used as input to Cyclum. Cyclum was run using a maximum of 5 linear dimensions and an encoder with 2 layers: the first containing 30 neurons, and the second containing 20 neurons.

*PCA*
For a given input transcript count matrix, genes with transcript pseudobulk proportions of $10^{-5}$ were kept. Transcript counts were then transformed by adding a pseudocount of 1 transcript, library size normalized, logged, and z-scored. The transformed count matrix was then transformed using PCA and the top 2 principal components were kept. Principal components were then scaled to [-1,1]. Using the scaled principal components, cell phase estimates were computed via the arctan2 function.
